# Dlk1 is a novel adrenocortical stem/progenitor cell marker that predicts malignancy in adrenocortical carcinoma

**DOI:** 10.1101/2024.08.22.609117

**Authors:** Katia Mariniello, James F.H. Pittaway, Barbara Altieri, Kleiton Silva Borges, Irene Hadjidemetriou, Claudio Ribeiro, Gerard Ruiz-Babot, Jiang A. Lim, Julie Foster, Julie Cleaver, Jane Sosabowski, Nafis Rahman, Milena Doroszko, Constanze Hantel, Sandra Sigala, Andrea Abate, Mariangela Tamburello, Katja Kiseljak-Vassiliades, Margaret Wierman, Laila Parvanta, Tarek E. Abdel-Aziz, Teng-Teng Chung, Aimee Di Marco, Fausto Palazzo, Celso E. Gomez-Sanchez, David R. Taylor, Oliver Rayner, Cristina L. Ronchi, Carles Gaston-Massuet, Silviu Sbiera, William M. Drake, Emanuel Rognoni, Matthias Kroiss, David T. Breault, Martin Fassnacht, Leonardo Guasti

**Author notes:** KM and JP contributed equally for this work (co-first authors).

## Abstract

Disruption of processes involved in tissue development and homeostatic self-renewal is increasingly implicated in cancer initiation, progression, and recurrence. The adrenal cortex is a dynamic tissue that undergoes life-long turnover. Here, using genetic fate mapping and murine adrenocortical carcinoma (ACC) models, we have identified a population of adrenocortical stem cells that express delta-like non-canonical Notch ligand 1 (DLK1). These cells are active during development, near dormant postnatally but are re-expressed in ACC. In a study of over 200 human ACC samples, we have shown DLK1 expression is ubiquitous and is an independent prognostic marker of recurrence-free survival. Paradoxically, despite its progenitor role, spatial transcriptomic analysis has identified DLK1 expressing cell populations to have increased steroidogenic potential in human ACC, a finding also observed in four human and one murine ACC cell lines. Finally, the cleavable DLK1 ectodomain is measurable in patients’ serum and can discriminate between ACC and other adrenal pathologies with high sensitivity and specificity to aid in diagnosis and follow-up of ACC patients. These data demonstrate a prognostic role for DLK1 in ACC, detail its hierarchical expression in homeostasis and oncogenic transformation and propose a role for its use as a biomarker in this malignancy.

**Graphical abstract:** 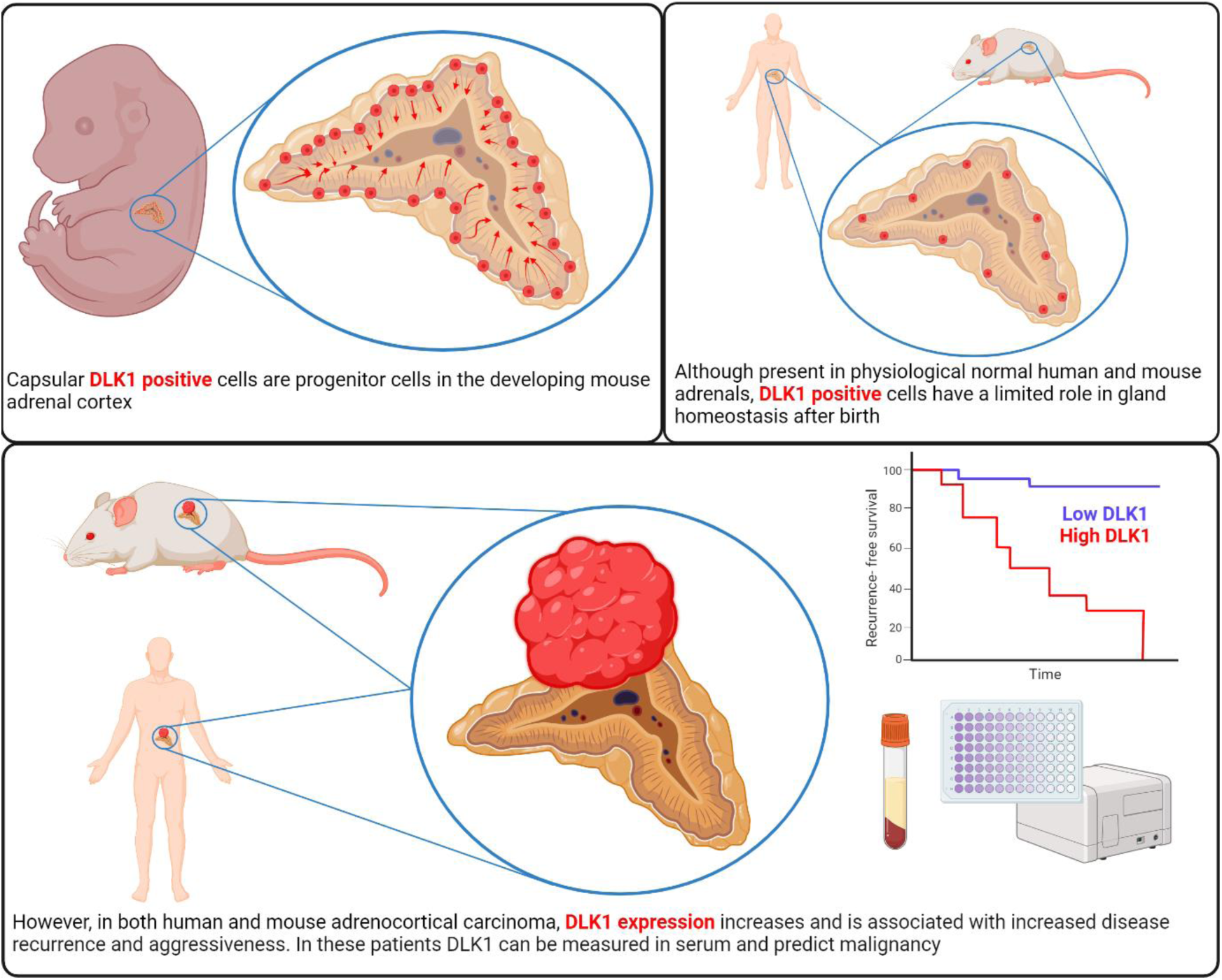

**Statement of significance:** This study presents DLK1 as a novel biomarker in ACC with opportunities for use in the diagnosis, prognosis and longitudinal follow up of patients. DLK1, a marker of adrenocortical stem cells, is re-expressed in ACC, is measurable in patients’ serum and is associated with increased malignancy.

## Introduction

Adrenocortical carcinoma (ACC) is a rare malignancy with a heterogeneous prognosis (1, 2). Complete early surgical resection offers the best chance of cure but frequently disease presents late and in the advanced stage five-year survival is <15% (3, 4). The only specifically approved medical treatment for ACC is the adrenolytic drug mitotane but it is poorly tolerated and often fails to prevent disease progression (5). Dysregulation of signaling pathways involved in the organogenesis and homeostasis of the adrenal cortex is implicated in the pathogenesis of ACC (6). Large pan-genomic analyses of ACC have identified alterations in Wnt/β-catenin, cAMP/PKA, and TP53 pathways as frequent molecular events in this malignancy (7–9). We have previously identified the protein delta-like non-canonical Notch ligand 1 (DLK1) to be co-expressed with sonic hedgehog in the adrenocortical progenitor niche (the “undifferentiated zone”) in the rat adrenal cortex (10). Moreover, we have shown that the human adrenal cortex remodels with age to generate clusters of relatively undifferentiated cells expressing DLK1 (11). DLK1 is a transmembrane protein with a cleavable ectodomain, belonging to the Notch/Delta/Serrate family. Paternally expressed, it is part of the group of imprinted genes located on chromosome band 14q32 in humans and 12qF1 in mice. During embryonic development, DLK1 is expressed at a high level in numerous human tissues, whereas in adults its expression is restricted to (neuro)endocrine tissues and other immature stem/progenitor cells, notably hepatoblasts. However, DLK1 expression is reported in a number of malignancies at a high frequency, where it is associated with worse survival outcomes (12). Indeed, we have previously demonstrated an increase in DLK1 expression in a small cohort of ACC compared with normal adrenal glands and benign aldosterone-producing adenomas (11). Overexpression of *DLK1* in ACC, compared to normal adrenals, was also recently demonstrated with single-nuclei sequencing (13). This, coupled with functional evidence that DLK1 inhibits differentiation, enhances cancer stemness and stimulates tumorigenesis has established its candidacy for further investigation in ACC (12).

Studies in ACC have traditionally been restricted through the rarity of the condition and paucity of appropriate preclinical models. Historically, murine models have failed to recapitulate the metastatic and aggressive nature of the disease. However, this has now been achieved through transgenic mouse models which carry genomic alterations seen in human ACC e.g. *p53*/*Rb* (14) and *p53*/*Ctnnb1* (15, 16). Additionally, several novel patient-derived ACC cell lines have been developed recently (17–20).

Here, we present a comprehensive characterization of DLK1 in normal adrenal physiology and ACC. Using genetic fate mapping, we demonstrate the contribution of DLK1 to adrenocortical development and self-renewal in mice. We draw upon the developments above to characterize DLK1 expression in both mouse models of adrenocortical tumorigenesis/carcinogenesis and patient-derived cell lines. Additionally, we present DLK1 expression data from two different patient cohorts, establishing the prognostic significance of DLK1 expression in ACC, highlighting a role for both tumor associated and secreted DLK1 as a novel biomarker in ACC, and exploring its function within tumors though spatial transcriptomic analysis.

## Results

### Dlk1 cells are capsular and cortical during embryonic development, and capsular only postnatally in mice

The developing adrenal gland showed widespread Dlk1 expression, with virtually all capsular cells displaying Dlk1 immunoreactivity up to embryonic day (E) 15.5 and maintaining high expression up to postnatal day (P) 0 (Figure 1 A-D and G). Clusters of subcapsular cortical cells expressing Dlk1 decreased during development with a few remaining at P0. High expression of Dlk1 in the medulla was detected at all stages analyzed. Postnatally, Dlk1 immunoreactivity was restricted to capsular cells and to the medulla (Figure 1 E-G). To determine the spatial relationship of Dlk1-expressing cells and subcapsular Axin2^+^ early adrenal progenitor cells (21), *Axin2^CreERT2/+^;Rosa^YFP/YFP^* mice (herein called *Axin2Cre*) were employed. These mice express the inducible Cre recombinase in Axin2^+^ cells, and Tamoxifen-induced recombination in the Rosa26 locus results in permanent labelling of Axin2-expressing cells and their progeny with Yellow Fluorescence Protein (YFP). After a 4-day chase (Figure 1 H), E19.5 cortical Axin2 and their early descendants were mostly Dlk1^-^, with 4-7% of cortical cells co-expressing Dlk1 and YFP (Figure 1 I-P). Postnatally, a 14-day chase (Figure 1 Q) showed the majority of YFP cells to be subcapsular, with a few migrating further into the Zona Fasciculata (ZF). Interestingly, YFP signal could be detected in 10-12% of postnatal capsular cells, and around one quarter of these were positive for Dlk1 in both males and females (Figure 1 R-X). Indeed, active β-catenin immunostaining was observed in capsular cells with immunohistochemistry (Figure S1 A and B). To further investigate the phenotype of Dlk1 capsular cells in postnatal adrenals, we employed a Platelet-derived Growth Factor Receptor α (PDGFRα^EGFP^) transgenic line, which expresses the histone H2B-enhanced Green Fluorescence Protein (eGFP) fusion protein from the endogenous *Pdgfrα* locus; Pdgfrα (CD140b) marks mesenchymal stem cells/fibroblastic cells (22). In these mice, a strong capsular GFP signal was detected (Figure S1C). While the majority of these cells were negative for Dlk1 expression, approximately 5% were double positive in both males and females (Figure S1 D-G). As expected, Pdgfrα^+^ cells were adjacent to, but distinct from, Cyp11b2-expressing Zona Glomerulosa (ZG) cells (Figure S1 F). Dlk1 cells were rarely positive for Ki-67 (<1% in the capsule and <5% in the subcapsular clusters during development, 0% in the postnatal capsule, Figure S1 H-M). *Gli1* expression in the capsule, different from Dlk1, remained high during development and throughout postnatal life (Figure S2). These data show that Dlk1 is both cortical and capsular during embryonic development and is restricted to the capsule postnatally. Additionally, previously unrecognized activity of the WNT pathway in some capsular cells, where Dlk1 is observed, raises the possibility of a functional interaction between the Dlk1 and the WNT pathway.

**Figure 1.**
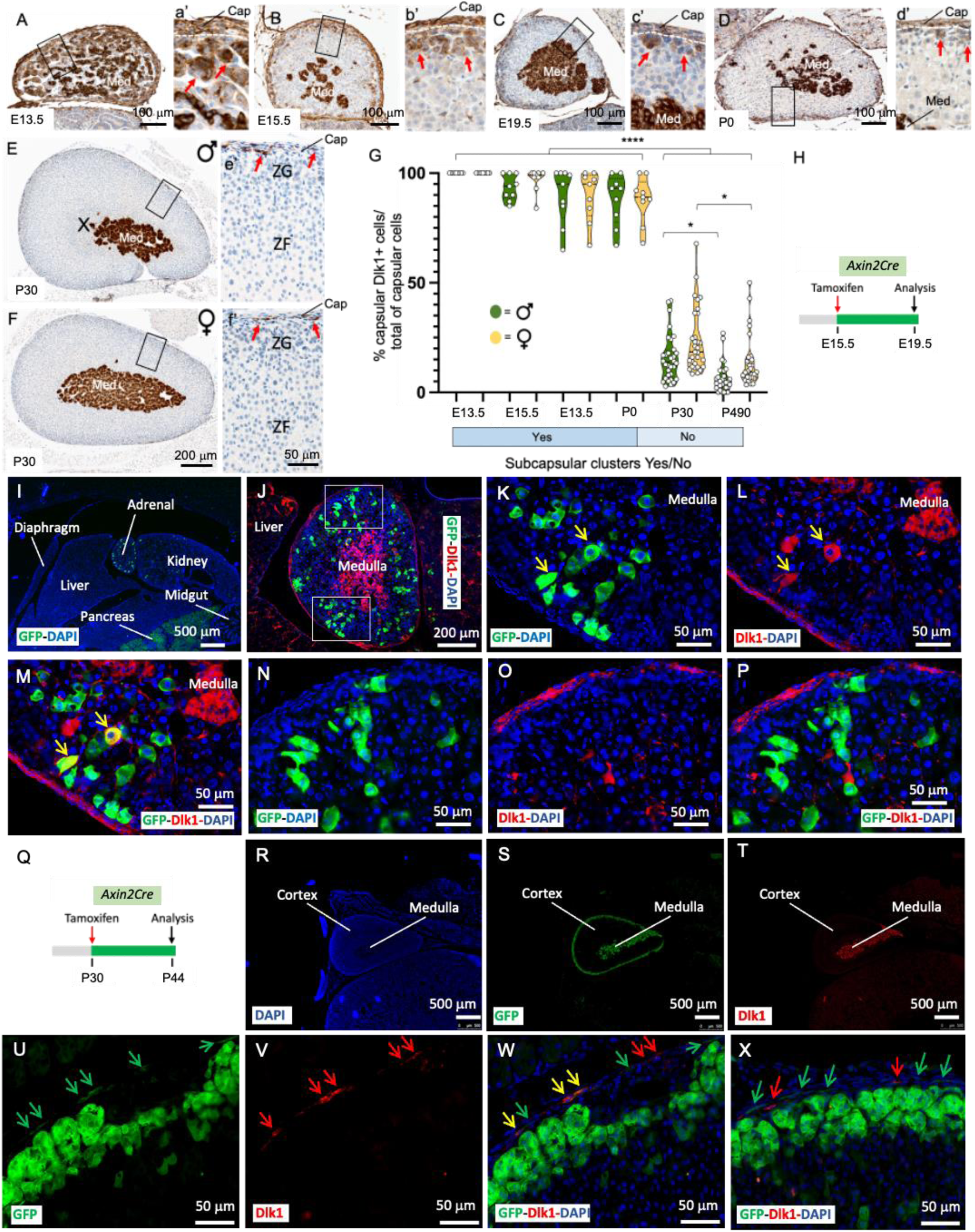
Embryonic and postnatal expression of Dlk1 in the mouse adrenal. A-d’) Immunohistochemical detection of Dlk1 in E13.5 (A and a’), E15.5 (B and b’), E19.5 (C and c’) adrenals showing expression in the capsule, cortex, and medulla. Subcapsular clusters of Dlk1^+^ cells (red arrows) decreased during development and were sparse at P0 (D and d’). E-f’) Immunohistochemical detection of Dlk1 in 4-weeks old female (E and e’) and male (F and f’) adrenals. G) Percentage of Dlk1^+^ cells in the capsule showed a dramatic reduction after birth, with a small non-significant trend of higher number of Dlk1^+^ cells in female mice. Cap, capsule; ZG, Zona Glomerulosa; ZF, Zona Fasciculata; Med, medulla; X-zone H) Schematic of tamoxifen treatment in *Axin-2Cre* mice during development. I-P) Localization of Axin-2^+^ cells and Axin-2 early progeny (4 days chase, green) relative to Dlk1 expression (red). Nuclei (DAPI) are in blue. Note the presence of occasional cortical GFP^+^/Dlk1^+^ cells (yellow arrows in K-M) and absence of GFP staining in the capsule, that instead is strongly Dlk1^+^. GFP^+^/Dlk1^+^ cells were TH^-^ and Sf1^+^ (not shown). Q) Schematic of tamoxifen treatment in postnatal *Axin-2Cre* mice. R-X) Localization of Axin2+ cells and Axin2 early progeny (14 days chase, green) relative to Dlk1 expression (red). Green arrows indicate capsular GFP^+^/Dlk1^-^ cells, yellow arrows indicate GFP^+^/Dlk1^+^ cells and red arrows Dlk1^+^/GFP^-^ cells. *p<0.05, ****p<0.0001.

### Dlk1 cells are adrenocortical stem cells active during development but dormant postnatally and upon postnatal ZF and ZG remodeling in mice

To assess whether Dlk1^+^ cells marked a population of adrenocortical progenitor cells, we employed genetic lineage tracing using inducible *Dlk1^CreERT2/+^;Rosa^tdTomato/+^* mice (herein called *Dlk1Cre*), where Dlk1^+^ cells and their progeny can be labelled with tdTomato (and labelled with an anti-Red Fluorescence Protein (RFP) antibody) upon tamoxifen injection. RFP expression was assessed with both immunofluorescence and immunohistochemistry. When dams were injected with tamoxifen at E12.5 and adrenals analyzed at both P10 and P38, clusters and columns of RFP^+^/Sf1^+^ cells could be observed spanning the whole width of the cortex (Figure 2 A-E). Dlk1 progeny significantly decreased at P38 compared to P10 and females showed a small non-significant trend of more Dlk1 progeny compared to males (Figure 2 F). Injection of tamoxifen of males and females at P0 and P30, followed by chases of 2 weeks, 1, 2 and 3 months did not show any cortical RFP^+^ cells (not shown), while longer chases (1 year and 2 years) showed occasional columns or cluster of RFP^+^ cells, representing <4% of the total cortical areas, in both males and females. In all cases, medullary cells were strongly RFP^+^, confirming effective recombination.

**Figure 2.**
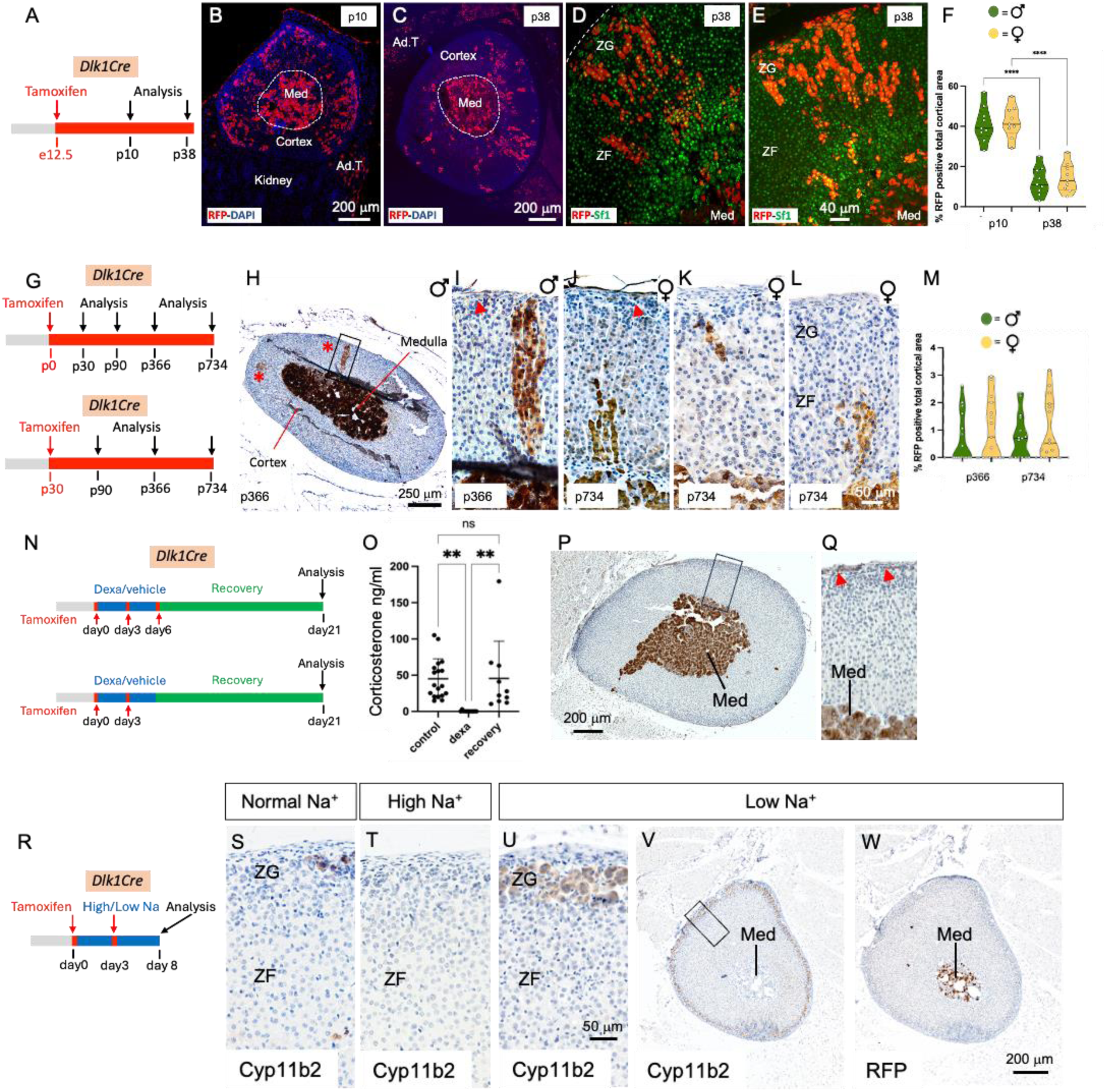
Dlk1 cells are adrenocortical progenitors during development, near-dormant postnatally and inactive upon adrenocortical remodeling. A) Schematic of tamoxifen induction of *Dlk1Cre* dams. B and C) Immunofluorescence detection of RFP^+^ cells at P10 (B) and P38 (C). Dlk1 progeny are positive for Sf1 expression in both males (D) and females (E). F) Time-course percentage of RFP^+^ cells in the cortex. G) Schematics of tamoxifen induction of *Dlk1Cre* in P0 and P30 mice. H-M) Examples of immunohistochemical detection of RFP^+^ cells after a year chase in male (H-J) and after a two-years chase in female (K and L) mice. Red asterisks in the panoramic panel H point to the occasional clusters and columns of RFP^+^ cells. These were TH^-^. Red arrows indicate capsular RFP^+^ cells. M) Time-course percentage of RFP^+^ cells in the cortex. N) Schematics of tamoxifen induction and dexamethasone treatment in P30 mice. O) Corticosterone levels measured before dexamethasone treatment, after dexamethasone treatment, and after ZF regeneration. P and Q) Immunohistochemical detection of RFP^+^ cells after ZF regeneration in *Dlk1Cre*. R) Schematic of tamoxifen induction of *Dlk1Cre* mice. S-V) Immunohistochemical detection of Cyp11B2 expression in mice fed with a normal (S), high sodium (Na^+^) (T), and low sodium (U, V) diet. W) Immunohistochemical detection of RFP^+^ cells after low sodium diet. The strong RFP staining in the medulla in B, C, H-L, P, Q, W suggests efficient recombination. Ad. T, adipose tissue; Med, medulla; ZG, Zona Glomerulosa; ZF, Zona Fasciculata. *p<0.05, **p<0.01, ***p<0.001, ****p<0.0001.

Capsular Gli1^+^ cells can be quickly programmed to replenish regenerating ZF cells following dexamethasone treatment (which induces ZF atrophy) (23). A 4-day dexamethasone regimen resulted in undetectable serum corticosterone in P30 mice, which recovered to pre-treatment levels 14 days after dexamethasone was stopped (Figure 2 O). In young (P30) and aged (P470) males and female *Dlk1Cre* mice, two different regimens of tamoxifen induction (Figure 2N, see Methods) resulted in no cortical RFP immunoreactivity during ZF regeneration (Figure 2 P and Q). Cortical RFP immunoreactivity was absent in corn oil (vehicle)- or tamoxifen-injected mice (not shown). In ZG remodeling, achieved through low sodium (ZG expansion) and high sodium (ZG regression) diets (Figure 2 R), neither male nor female P50/70 tamoxifen-injected *Dlk1Cre* mice showed RFP immunoreactivity in the ZG (or elsewhere in the cortex), despite profound changes in Cyp11b2 expression (Figure 2 S-W). All mice (ZF regeneration and ZG remodeling) had strong RFP staining in the medulla, confirming effective recombination. Taken together, these results demonstrate that Dlk1^+^ cells represent a novel population of adrenocortical progenitor cells in mice, which are active during embryonic development and near-dormant postnatally. Neither postnatal short-term ZG nor ZF remodeling activate Dlk1 capsular cells to generate cortical progeny.

### Adrenal subcapsular hyperplasia in mice is not derived from Dlk1^+^ cells

An abundance of capsular-like cells is a pathognomic feature in adrenal subcapsular hyperplasia (SH), a histological hallmark in mouse adrenals which occurs spontaneously mostly in aged females but also in certain strains/transgenics after gonadectomy (GDX). In the latter, the ensuing excess of gonadotrophins reprogram adrenal cells to a gonadal phenotype (i.e. activation of Gata4 transcription factor), and SH foci are thought to represent a morphologic continuum progressing to adrenocortical tumors (24, 25). SH contains cells formed by large sex steroid-secreting, lipid-laden B cells interspersed in aggregates of lipid-depleted, spindeloid, capsular-like, Sf1^-^ A cells, and can expand deep into the cortical parenchyma. As Dlk1 is expressed in the capsule (Figure 1) and marks adrenocortical progenitors (Figure 2), we hypothesized that SH and SH-derived adrenal tumors in these models may be enriched in and even derived from Dlk1^+^ cells. We employed GDX DBA/2J (26), GDX inhibin α subunit promoter (Inhα)/Simian virus 40 T-antigen (Tag) (27) and aged C57BL/6J for Dlk1 expression, and *Dlk1Cre* mice for lineage tracing. In GDX DBA/2J and Inhα/Tag, Dlk1 was not detectable in adrenal SH nor in the established tumors which instead strongly expressed Gata4 (Figure S3). As Dlk1 might still be the cell of origin of SH, but turning its expression off, we assessed expression of RFP in SH foci in older *Dlk1Cre* mice (1 and 2 years) which were injected with Tamoxifen at P0 or P30. Here, SH cells were found to be RFP negative in both males and females (Figure S4). Of note, SH A cells were PDGFRα^+^ and *Gli1^+^* (Figure S4), suggesting the contribution of different capsular fibroblast-like populations to SH formation. Altogether, these data showed that GDX-induced SH and SH-derived adrenal tumors are not enriched in Dlk1-expressing cells and that spontaneous SH foci in aged mice are not enriched in nor derived from Dlk1-expressing cells.

### Dlk1 is re-expressed in an autochthonous mouse model of ACC

Recently, a novel mouse model in which concomitant inactivation of *Trp53* and activation of *Ctnnb1* driven by the Aldosterone Synthase (AS)/CYP11B2 promoter (BPCre^AS/+^, herein called *BPCre*), was shown to recapitulate human ACC formation with high penetrance (15). The incidence of tumor formation and malignancy increases with age and by 12 months, all mice had generated adrenal tumors (86% ACC). We analyzed 23 tumor samples from 17 mice (9 female, Figure 3, at different ages (and therefore different stages of ACC development). Random sections were processed for both Dlk1 IHC and *Dlk1* RNAscope, showing an identical expression pattern (Figure S5A and B). There was low/no expression of Dlk1 in benign tumors, moderate expression in localized ACC and higher expression in metastatic disease, both in the primary tumors and in lung metastases. Generally, Dlk1 showed both a diffuse and a clustered (if not outright clonal), pattern of expression (Figure 3 A-F). When looking at all ACC samples in the cohort, there was a clear stepwise progression of Dlk1 expression with disease severity, localized ACC < Metastatic ACC < Metastases. The mean expression in these groups was different (F=10.89, p=0.0014**) (Figure 3G). *Post hoc* analyses also confirmed that the mean expression in metastases was higher than in non-metastatic ACC primary tumors (165.7 ± 57.70 vs 34.70 ± 43.27, adj. p=0.0010***). There was no difference in Dlk1 expression between sex of the mice (male 80 ± 38.21 vs female 66.07 ± 54.58, p=0.5563) (not shown). There was a positive correlation between Dlk1 expression levels and age of the mice (r=0.6007, p=0.0024**) (Figure 3H). Given aged mice tend to display more aggressive tumors, this finding is in keeping with the fact that Dlk1 expression is higher in more advanced disease in this model. These results indicate that in the *BPCre* model, Dlk1, rather than marking the cell of origin of ACC, is re-expressed in ACC, potentially conferring characteristics of undifferentiated cancer cells.

**Figure 3.**
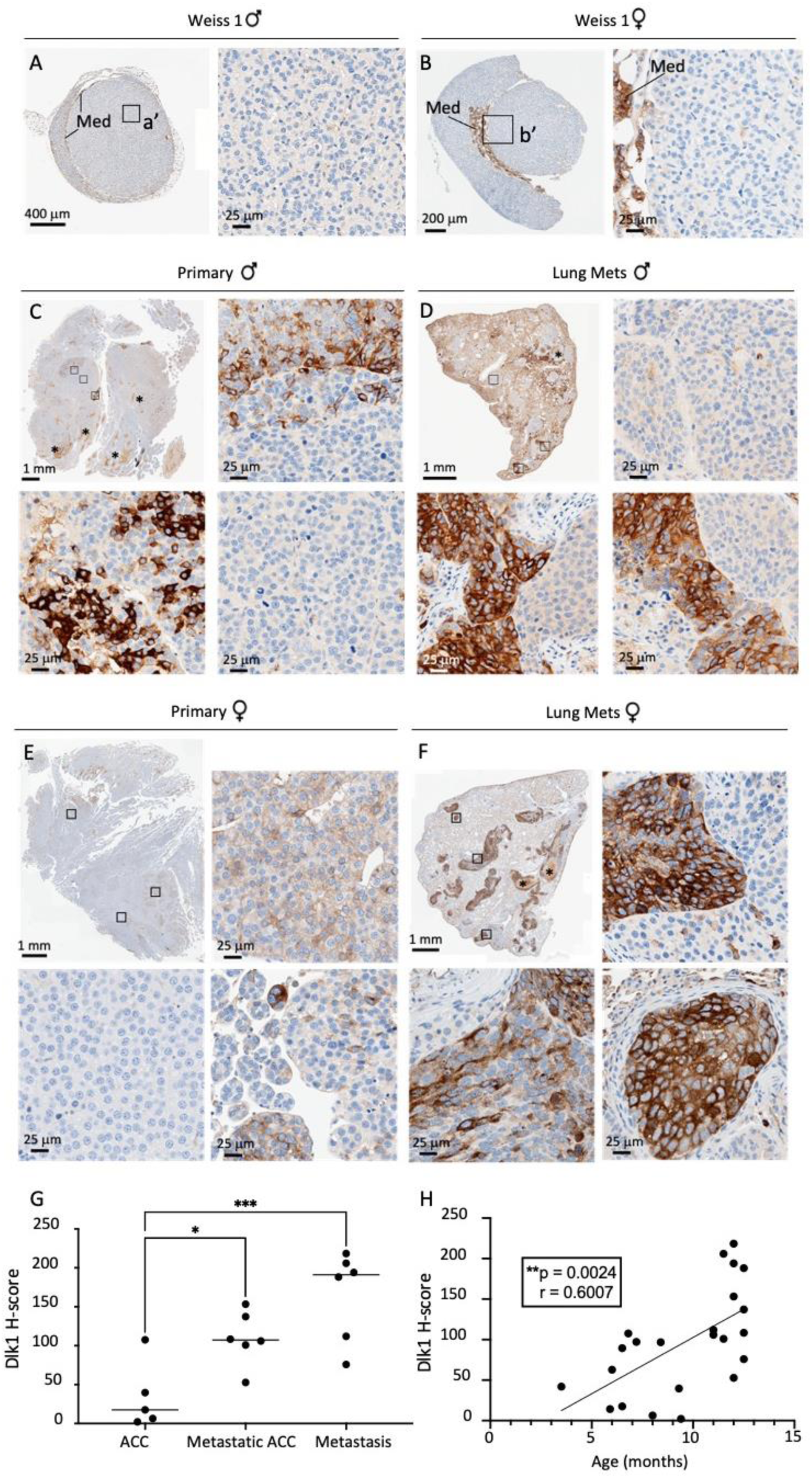
Dlk1 is re-expressed in a murine model of ACC and exhibits intratumoral heterogeneity. A-F) Immunohistochemical detection of Dlk1 expression in ACC from *BPCre* male (A) and female (B) mice with low Weiss Score, in metastatic ACC (C, primary male; E, primary female) and in lungs metastasis (Mets, D, males; F, females). Note the higher expression of Dlk1 in metastatic ACC and lungs metastasis. Dlk1 expression is mostly clonal with different foci that express varying levels of Dlk1 or be negative for Dlk1 expression. G) Dlk1 expression increases in a stepwise manner from non-metastatic primary ACC to metastatic primary ACC and metastatic lesions. Horizontal lines represent group means. H) Dlk1 expression positively correlates with age, which in turn increases with disease malignancy. Each dot represents an individual tumor. *p<0.05, **p<0.01, ***p<0.001.

### DLK1 expression is higher in human ACC than other pathologies and normal adrenal tissue

We previously showed in a small sample cohort DLK1 expression is significantly higher in ACC compared to normal adrenal tissue and aldosterone-producing adenomas (APA) (11). Moreover, analysis of PanCancer RNA-seq data across 29 cancer histotypes showed the highest expression of *DLK1* was in ACC and pheochromocytomas (28). To corroborate these findings, a prospective discovery cohort was established in London. 73 consecutive patients (26 male) undergoing adrenalectomy for suspected ACC or functioning benign adrenal pathology were recruited (Table S1). DLK1 expression was higher in ACC (n=12) than other benign pathologies (n=29) and normal adrenal samples (n=16) (F=5.937, p=0.0005***). *Post hoc* analyses revealed the mean H-score in ACC (115.4 ± 89.2) was higher than each individual group when assessed with multiple comparisons and adjusted p values (adrenal adenoma 54.27 ± 56.88, p=0.032*; normal adjacent 29.06 ± 39.77, p=0.0020**; other benign 10.34 ± 19.38, p=0.0092**; other malignant 1.806 ± 3.032, p=0.0040**) (Figure S6 A). As for *BPCre*, random ACC sections were also processed in parallel for *Dlk1* RNAScope, showing again an identical pattern of expression (Figure S4 C and D).

These findings were validated in a larger cohort of ACC tumor samples at the University Hospital Würzburg (Germany). The cohort consisted of 159 patients (53 male). From these, a total of 178 ACC tissue sections were studied. 15 patients had more than one tissue section in the cohort (11 with primary tumors and secondary disease and 4 with primary tumors and two specimens of secondary disease). The clinical and histopathological details of the cohort are reported in Table 1. DLK1 expression was seen in every tissue sample studied. There was a wide range of expression across the cohort (H-score 10 – 244, median 131.5) (Figure 4 A and B). Commonly, the expression pattern of DLK1 in the tumor sections was heterogenous with apparent clones of DLK1 positive cells, similarly to *BPCre* mice, and other areas of tumor parenchyma which were DLK1 low/negative (Figure 4 C and D). DLK1 was not found to be expressed in the connective tissue or in the associated vasculature. There was no correlation between DLK1 expression and patient age (r=-0.03216, p=0.6717) or sex (female H-score 117.0 ± 56.39, male H-score 134.6 ± 59.76, p =0.0709) (Figure S6 B and C). Tumor size also had no bearing on the level of DLK1 expression both in primary tumors (r=0.02189, p=0.8070) and secondary disease (r=-0.06547, p=0.6842) (Figure S6 D and E).

**Figure 4.**
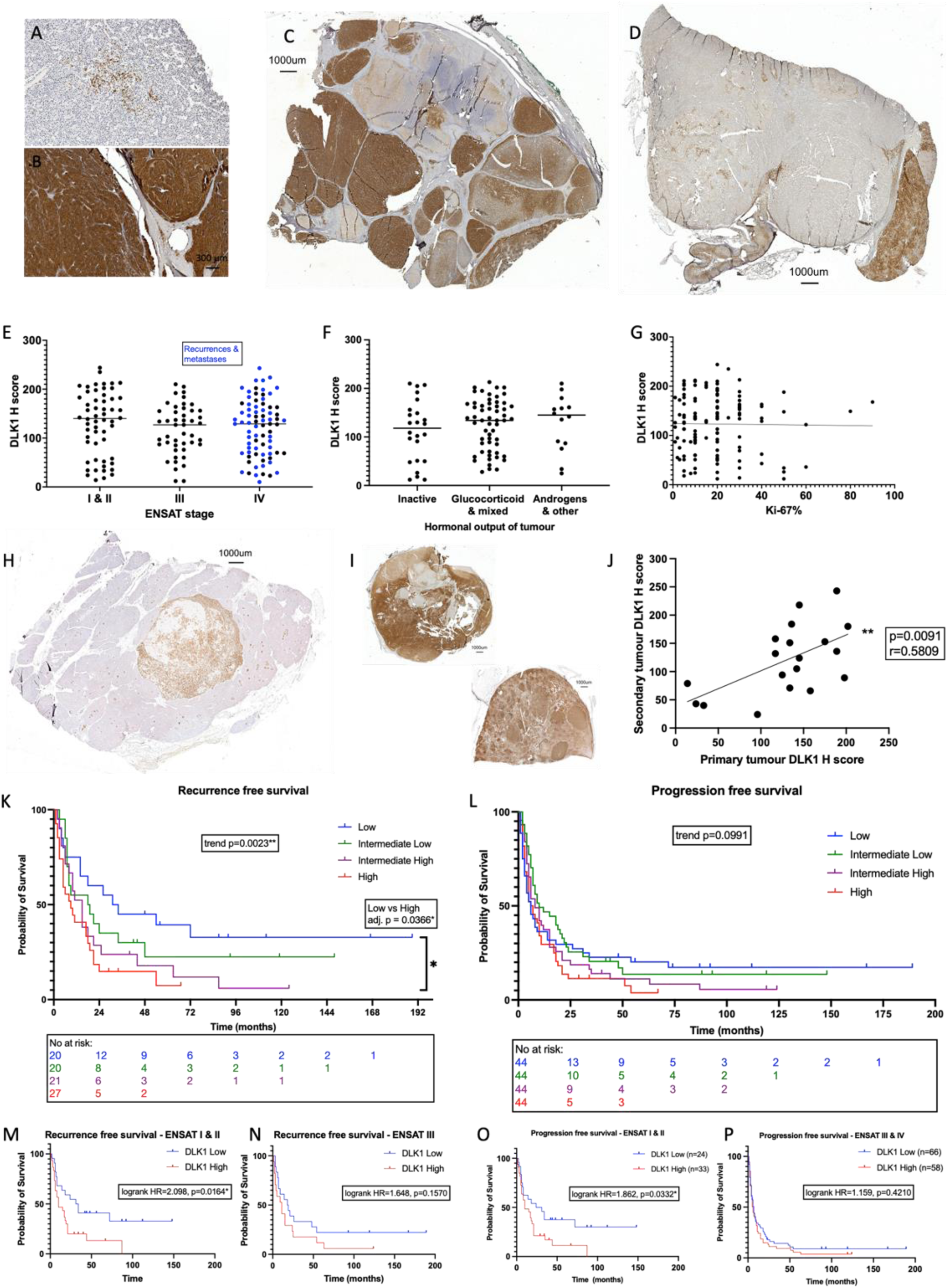
DLK1 expression is ubiquitous in human ACC, consistent in metastatic disease and increases risk of disease recurrence and progression. A-G) The range of expression across the cohort can be seen with few positive cells in the tumor parenchyma (A) dense, intense staining throughout (B). C and D: panoramic sections illustrating heterogenous DLK1 expression in individual tumor samples. DLK1 expression is unrelated to ENSAT stage (E), hormonal output (F) or Ki-67% (G). H-J) DLK1 expression shown in a liver metastasis (H). DLK1 expression is positively correlated in primary and secondary disease in the same patients (I and J). Each dot represents a secondary tumor. K-L): Kaplan-Meier curves showing increased disease recurrence (K) and a trend towards increased disease progression (L) with higher DLK1 expression levels. This effect is more pronounced in ENSAT stage I & II disease (M, O) than in ENSAT stage III & IV disease (N, P). *p<0.05, **p<0.01.

**Table 1.** Descriptive characteristics of tissue validation cohort (Würzburg) * Weiss 2 ACC originally classified as ACA but then progressed

|  |  | n | Median | Range |
| --- | --- | --- | --- | --- |
| **Sex** | Male | 53 |  |  |
|  | Female | 105 |  |  |
|  | Total | 159 |  |  |
| **Age at diagnosis (years)** |  |  | 47 | (16-80) |
| **ENSAT Tumor Stage** | I | 9 |  |  |
|  | II | 79 |  |  |
|  | III | 51 |  |  |
|  | IV | 39 |  |  |
| **Tumor entity** | Primary | 131 |  |  |
|  | Local recurrence | 27 |  |  |
|  | Metastasis | 20 |  |  |
|  | Total | 178 |  |  |
| **Tumor size (cm)** | Primary |  | 10.3 | (3-24) |
|  | Local recurrence |  | 3.8 | (1.1-9.4) |
|  | Metastasis |  | 1.7 | (0.7-5.5) |
|  | Total |  | 9 | (0.7-24) |
| **Hormone secretion** | Cortisol (alone or mixed) | 72 |  |  |
|  | Other | 16 |  |  |
|  | Inactive | 29 |  |  |
|  | Unknown | 61 |  |  |
| **Resection Status** | 0 | 86 |  |  |
|  | 1 | 14 |  |  |
|  | 2 | 42 |  |  |
|  | X | 36 |  |  |
| **Weiss Score** |  |  | 6 | (2*-10) |
| **Ki-67%** |  |  | 20 | (1-90) |

DLK1 expression was consistent across the different ENSAT tumor stages at presentation (ENSAT I&II 130.7 ± 64.97, ENSAT III 117.4 ± 50.30, ENSAT IV 121.4 ± 57.67, ANOVA F=0.7307, p=0.4830) (Figure 4E). There was no difference in DLK1 expression in the hormonal activity of tumors (inactive 111.8 ± 63.38, active 126.4 ± 52.97, p=0.2695). This was also shown when looking at different categories of associated hormonal excess, glucocorticoids or other (ANOVA F=0.6273, p=0.5363) (Figure 4F). Additionally, DLK1 expression was unrelated to Weiss score (r=0.03056, p=0.7469) or Ki-67% (r=-0.01395, p=0.8778) (Figure 4G).

DLK1 expression was present in recurrent disease and could clearly identify metastases from background tissue (Figure 4 H and I). 19 secondary disease specimens were available in patients whose primary tumors were included in the study. The DLK1 expression level seen in the secondary disease demonstrated a positive correlation with the expression level in the primary tumors (r=0.5809, p=0.0091**) (Figure 4 J).

### Higher levels of DLK1 expression were associated with an increased risk of disease recurrence in patients with ACC

In the 88 primary tumors of the German cohort who were disease free after surgery, higher levels of DLK1 expression were associated with a doubling in the risk of disease recurrence compared with lower DLK1 levels (median recurrence-free survival high DLK1 10.5 months vs 22.5 months in low DLK1, HR 1.979 95% CI 1.218 - 3.216, p=0.0059**). This was more pronounced in ENSAT stage I & II (n=52, median recurrence-free survival high DLK1 10 months vs 32.5 months in low DLK1, HR 2.098 95% CI 1.127 – 3.903, p=0.0164*) than in ENSAT stage III & IV disease (n=36, median recurrence-free survival high DLK1 11 months vs 18.5 months in low DLK1, HR 1.648 95% CI 0.7961 – 3.412, p=0.1570) (Figure 4 M and N). Further to this, when categorizing DLK1 expression levels in quartiles (based on median and interquartile range values), higher DLK1 levels were associated with stepwise increased risk of recurrence (median recurrence-free survival low DLK1 32.5 months, low-intermediate DLK1 18.5 months, high-intermediate DLK1 15 months and high DLK1 9 months). This was significant both by log-rank test for trend across the four groups (χ^2^=9.263, p=0.0023**) and when comparing the high vs low DLK1 expression groups directly (adjusted p=0.0185*) (Figure 4 K). Cox regression analysis revealed that the only statistically significant variables associated with recurrence-free survival in this cohort were Ki-67% and DLK1 expression (Table 2). Multivariate analysis with these two variables revealed that they both were independent risk factors of disease recurrence (Table 2). Higher DLK1 levels were the strongest predictor of disease recurrence in this model.

**Table 2.**
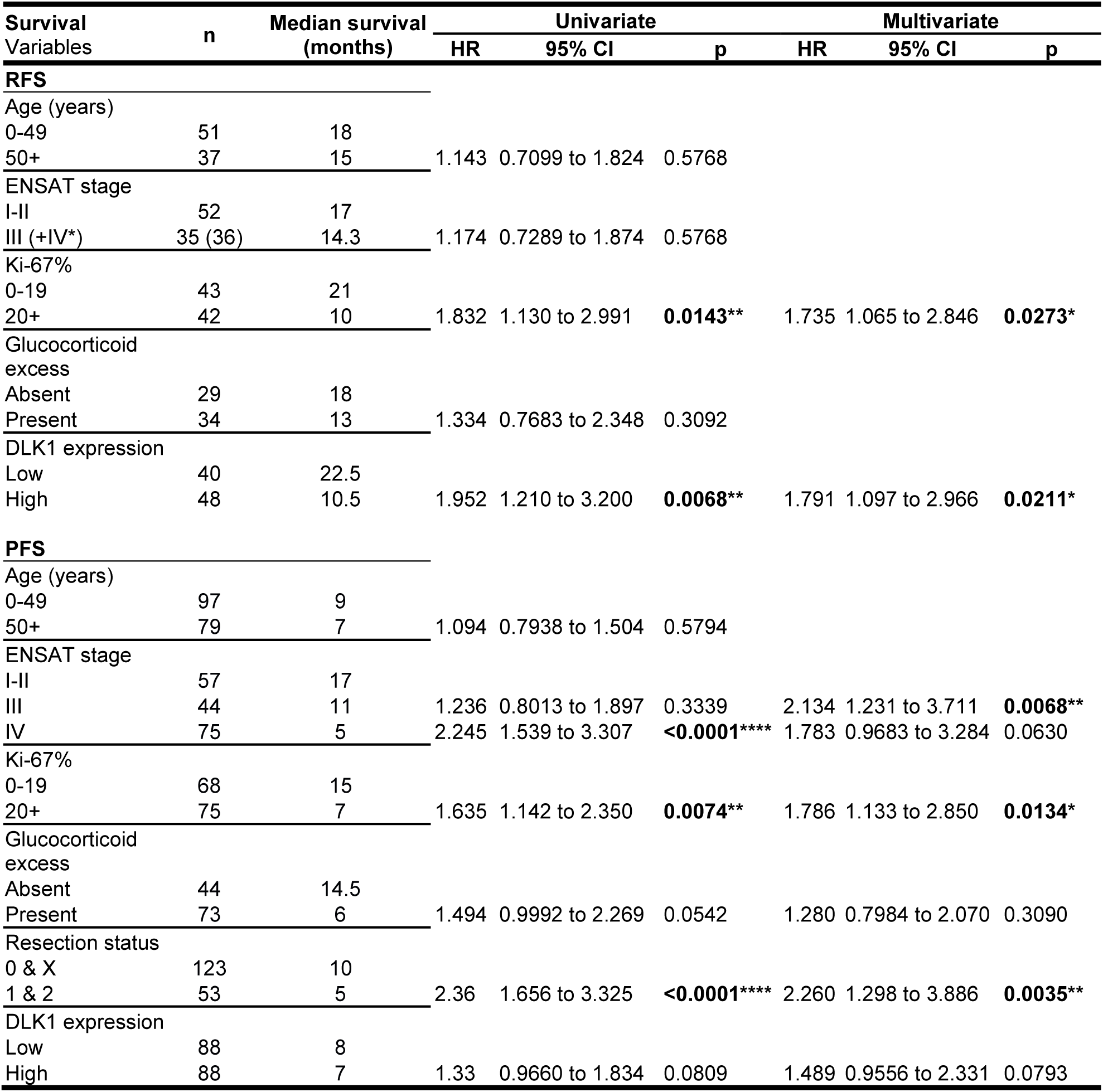
Cox regression analyses of recurrence-free and progression-free survival in the validation cohort (Würzburg) RFS – recurrence-free survival, PFS – progression-free survival

Higher DLK1 levels were associated with a trend towards increased risk of disease progression (n=176, median progression-free survival high DLK1 7 months vs 8 months in low DLK1, HR 1.311 95% CI 0.9538 – 1.801, p=0.0801). This trend was also seen when assessing DLK1 expression in quartiles (log-rank test for trend χ^2^=2.72, p=0.0991) (Figure 4L). In the entire cohort, the trend of higher DLK1 expression being associated with decreased survival appeared to be more prominent after approximately 24 months and was not clearly reflected in the median survival times (median survival low DLK1 6 months, low-intermediate DLK1 10 months, high-intermediate DLK1 8 months and high DLK1 6.5 months). The disease progression risk of higher DLK1 expression was more pronounced in ENSAT stage I & II disease (n=57, median progression-free survival high DLK1 10 months vs 27.5 months in low DLK1, HR 1.863 95% CI 1.038 – 3.340, p=0.0332*) (Figure 4 O). In ENSAT stage III & IV groups median progression-free survival was comparable (n=121, high DLK1 6 months vs 7 months low DLK1, HR 1.159 95% CI 0.7931 – 1.694, p=0.4210) (Figure 4 P). Cox regression analysis revealed univariate influencers of worse progression-free survival were ENSAT stage, Ki-67% and resection status (Table 2). DLK1 expression level high vs low did not reach statistical significance in univariate analysis (HR 1.33 95%CI 0.9660 – 1.834, p=0.0809). Multivariate analysis, including all variables from univariate analysis with p value of <0.2, revealed that higher DLK1 levels were associated with a trend towards independence as a risk factor for progression-free survival (HR 1.489 95%CI 0.9556 – 2.331, p=0.0793). ENSAT stage, Ki-67% and resection status remained independent risk factors for progression (Table 2).

DLK1 expression was not associated with overall survival (n=131, median survival high DLK1 46 months vs 47 months in low DLK1, p=0.7161) (Figure S6 F). In patients with ENSAT stage I & II disease, lower DLK1 levels were associated with a non-significant trend towards longer overall survival (n=56, median survival high DLK1 48 months vs 91 months in low DLK1, HR 1.353 95%CI 0.6374 - 2.873, p=0.4310) (Figure S6 G). This trend was much less pronounced in ENSAT III & IV group (n=76, median survival high DLK1 26 months vs 36 months in low DLK1, HR 1.068 95%CI 0.6312 - 1.808, p=0.8056) (Figure S6 H). Cox regression modelling was performed to look more closely at the factors influencing overall survival (Table S2). Univariate analysis revealed that in this cohort, significant influencers were ENSAT stage, Ki-67% and resection status. In this model DLK1 expression levels did not have any bearing on overall survival (HR 1.081 95%CI 0.7061 – 1.656, p=0.7188). Multivariate analysis including the statistically significant variables in univariate analysis and DLK1 expression, found that the only independent effector of overall survival in this cohort is resection status.

### Serum Dlk1 levels are elevated in mouse models of adrenocortical carcinogenesis and are predictors of malignancy in humans

Dlk1 can be cleaved in the juxtamembrane region and the bioactive ectodomain released into the extracellular space (29). In mice and humans, serum Dlk1 levels are usually very low, except in the later stages of pregnancy, where mothers have high levels which, in mice, have been shown to be fetal in origin (30). Several studies have shown that DLK1 levels are measurable in blood of patients with cancers known to express DLK1 (31, 32). Recently it has been shown that serum Dlk1 levels correlate with tumor size in murine ovarian cancer (33). We have cloned two isoforms from both the human adrenal and H295R cells, corresponding to the known full-length and short isoforms, the latter lacking a small extracellular juxtamembrane region. A preponderance of expression of the longer isoform was confirmed in the human adrenal, H295R, and human ACCs (Figure 5 A). These isoforms were cloned in frame with an HA-tag at the N-terminus and a FLAG-tag at the C-terminus and expressed in HEK-293 cells (a cell line not expressing endogenous DLK1) (Figure 5 B). Medium from these cells was found to contain the cleaved ectodomain from both the short and long isoform by ELISA (Figure 5 D). We then assessed endogenous serum Dlk1 levels in BPCre mice and in a subcutaneous tumor mouse model injected with a BPCre tumor-derived cell line, BCH-ACC3A (16) (Figure 5 G). In both, serum Dlk1 levels were significantly higher compared to aged-matched controls, and a positive correlation between tumor size and serum levels was observed (Figure 5 E and F). Additionally, we injected H295R cells subcutaneously in Nu/Nu female mice (Figure 5 I and J) and allow tumors to grow to different sizes before blood collection. Serum Dlk1 levels, assayed with a human specific DLK1 ELISA, showed positive correlation with tumor size (Figure 5 H). These results strongly suggest that in mouse models of ACC, serum Dlk1 is shed from tumors.

**Figure 5.**
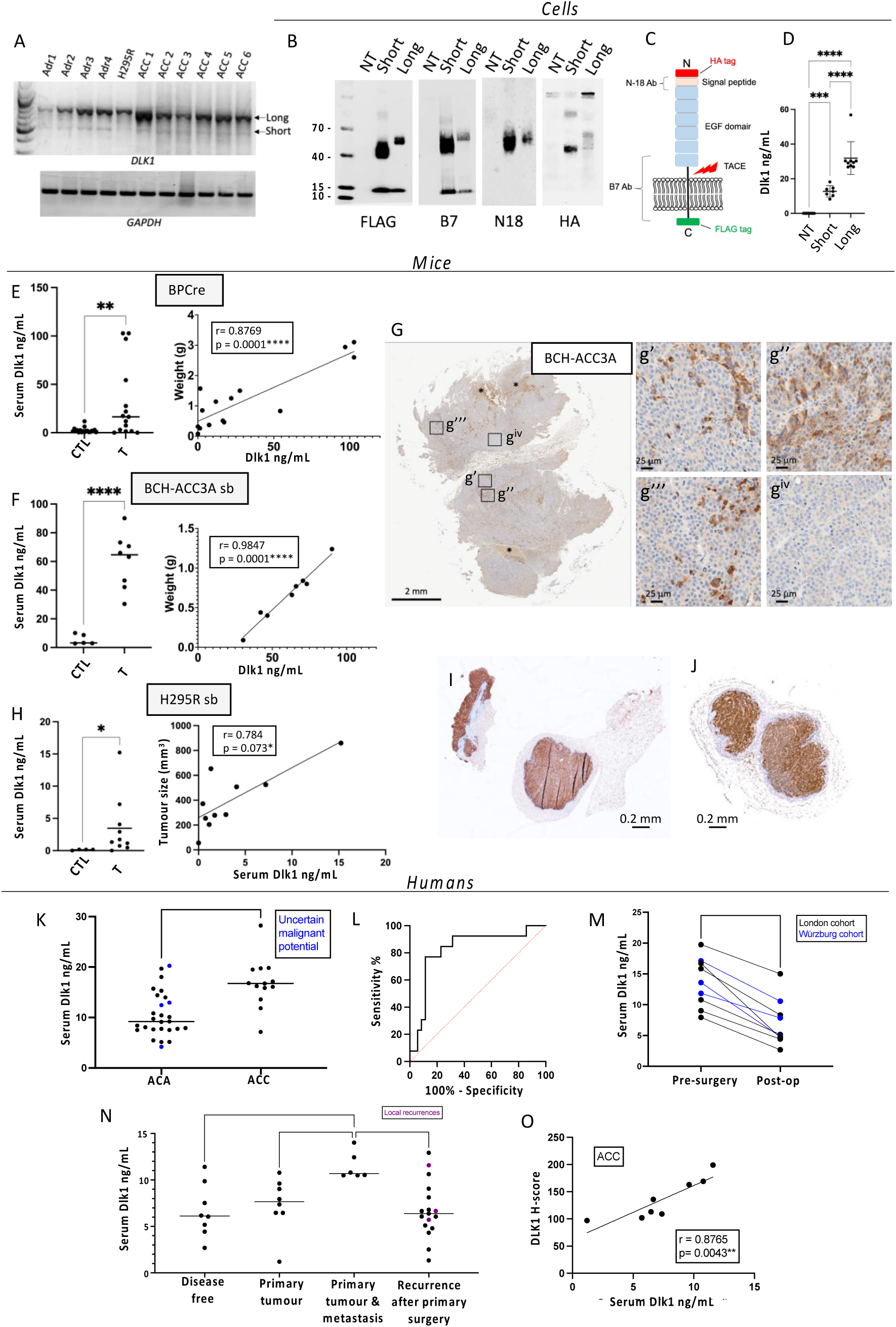
Serum DLK1 is novel biomarker in ACC that predicts malignancy. A) RT-PCR analysis for the expression of the full-length (Long) *DLK1*, short *DLK1* and *GAPDH* in normal human adrenals (Adr), H295R cells and six human ACCs. B) Western Blotting analysis of the two DLK1 isoforms (HA and FLAG tagged) transfected into HEK293 cells, using anti-FLAG (tag at the C-terminus), anti B7 (targeting aa 266-383), anti N18 (targeting the N-terminus region) and HA (tag at the N-terminus) antibodies. NT, non-transfected HEK293 cells. The lower band with an apparent molecular weight of 12 kDa, recognized only by the anti-FLAG and anti B7 antibodies, represents the membrane tethered DLK1 post cleavage mediated by Tumor necrosis Alpha Converting Enzyme (TACE). C) Schematic of DLK1 structure indicating regions targeted by antibodies. D) Levels of human DLK1 ectodomain in the medium. E) Levels of mouse Dlk1 ectodomain in the serum of age-matched controls (CTL) and *BPCre* mice (T) (left panel), and correlation with tumor weight (right panel). F) Levels of mouse Dlk1 ectodomain in the serum of aged-matched (CTL) and mice injected subcutaneously with BCH-ACC3A cells (T) (left panel), and correlation with tumor weight (right panel). G) Immunohistochemical detection of Dlk1 in tumors of mice injected subcutaneously with BCH-ACC3A cells, showing different levels of Dlk1 expression. H) Human DLK1 is not detected in serum samples from non-injected Nu-Nu mice using human-specific DLK1 ELISA (CTL), while a significant signal is obtained from sera from mice injected with H295R cells (T) (left panel). The correlation with tumor size is reported on the right panel. I-J) Examples of Dlk1 expression in tumors retrieved from Nu-Nu mice injected with H295R. K-L) *London cohort*. Pre-operative serum DLK1 levels are higher in those with ACC than ACA (K) and can predict malignancy as seen on ROC curve (L). M) In all patients studied, a significant fall in serum DLK1 levels was detected after resection of the primary ACC. N-O) *Würzburg cohort*. N) Serum DLK1 levels are higher in those presenting with ENSAT stage IV disease than other groups. Each dot is a blood sample relating to an individual patient. Purple dots represent local recurrences and horizontal lines represent group mean values. O) Serum DLK1 levels positively correlate with tissue expression (H-score) in the same patients with ACC. *p<0.05, **p<0.01, ***p<0.001, ****p<0.0001.

In humans, pre-operative serum DLK1 levels were measured in the London prospective discovery cohort (n=73). Descriptive characteristics for the sample cohort used in this study are shown in Table S1. Serum DLK1 levels were significantly higher in ACC (16.81 ± 4.876ng/mL) than in benign adrenocortical adenomas (10.54 ± 4.417ng/mL, p=0.0002***) (Figure 5 K). Using all pre-operative values, a receiver operator characteristic (ROC) curve showed that serum DLK1 levels were able to predict the diagnosis of ACC in this cohort (AUC 0.8242 ± 0.07214, p=0.0006***) (Figure 5 L). Serum DLK1 levels >15.77ng/mL predict ACC diagnosis in this cohort with a sensitivity 77% and specificity 89%. Similar to the tissue expression findings, DLK1 serum levels had no significant correlations with age, tumor size or Ki-67% (Figure S7 A-C).

The findings from the London serum cohort were validated in a separate cohort from Würzburg (n=25). All patients had a diagnosis of ACC and were characterized by different states of disease (Table S3). Patients presenting initially with ENSAT stage IV disease had higher serum DLK1 levels than patients with disease recurrence following primary surgery (11.46 ± 1.459ng/mL vs 6.749 ± 3.016ng/mL, adj. p=0.0058**), disease free patients (6.666 ± 2.855ng/mL, adj. p=0.0154*) and isolated primary tumors (11.46 ± 1.459ng/mL vs 7.357 ± 2.913ng/mL, adj. p=0.0469*) (Figure 5 N). Serum DLK1 levels did not correlate with tumor size, hormonal secretion of tumor, or Ki-67% in the validation cohort (Figure S7 F-H).

In a small number of ACC patients, post-operative bloods samples were also taken (n=9). The median time after surgery for these samples was 44 days (range 11-122). Due to smaller numbers and nature of analysis (paired t-test), samples were collated from both the London (n=6) and Würzburg (n=3) cohorts. All post-operative samples were taken at a time the patient was understood to be free of disease. Analysis showed that there was a significant decrease in serum DLK1 levels post-surgery (mean decrease -6.568 ± 2.565ng/mL, p<0.0001****) (Figure 5 M), suggesting serum DLK1 in pre-operative patients is shed from ACCs.

For patients in whom tissue and pre-operative serum was available in the London cohort, there was a significant positive correlation between DLK1 H-score in tissue and serum DLK1 levels (r=0.5131, p=0.0012**) (Figure S7 D). This was validated also considering the Würzburg cohort separately (r=0.8765, p=0.0043**) (Figure 5 O).

### DLK1^+^ cells are endowed with both enhanced steroidogenic potential and clonogenicity

To characterize in detail the transcriptomic differences between DLK1^+^ and DLK1^-^ areas within the ACC tumor parenchyma, GeoMX spatial whole-transcriptome profiling was performed by selecting 60 DLK1^+^ and DLK1^-^ Regions of Interest (ROI) within four human ACCs. Figure S6 illustrates a detailed schematic diagram of our protocol. Principle component analysis revealed the transcriptomic signatures of the positive and negative areas from different tumors clustered together in distinct groups. 1072 significant differentially expressed genes were identified between the positive and negative groups (adj. p<0.01). Unsupervised heatmap clustering of the differentially expressed genes revealed the transcriptomes of the positive and negative areas were distinct (Figure 6 A-B). By applying a fold change cut off of >/<2, there were 10 upregulated and 17 downregulated genes identified in the positive areas compared to the negative areas (Figure 6 B). Surprisingly, out of the 9 (excluding DLK1 itself) upregulated genes in the positive tumor areas, 5 are involved in the synthesis of cholesterol (EBP, DHCR7, DHCR24, MSMO1) and fatty acids (FADS2). The others are involved in steroidogenic pathway (CYP17A1), vesicular and cholesterol binding protein (SYP) and the genes for cathepsin (CTSA) and clusterin (CLU). The downregulated genes included pro-apoptotic genes (BNIP3, BNIP3L, NR4A1) and transcriptional regulators of differentiation (EGR1, FOS, JUN) (Figure 6 B). Interestingly, independent studies have demonstrated that FOS is a negative regulator of SF1 transcriptional activity (34) and CYP17A1 expression (35, 36).

**Figure 6.**
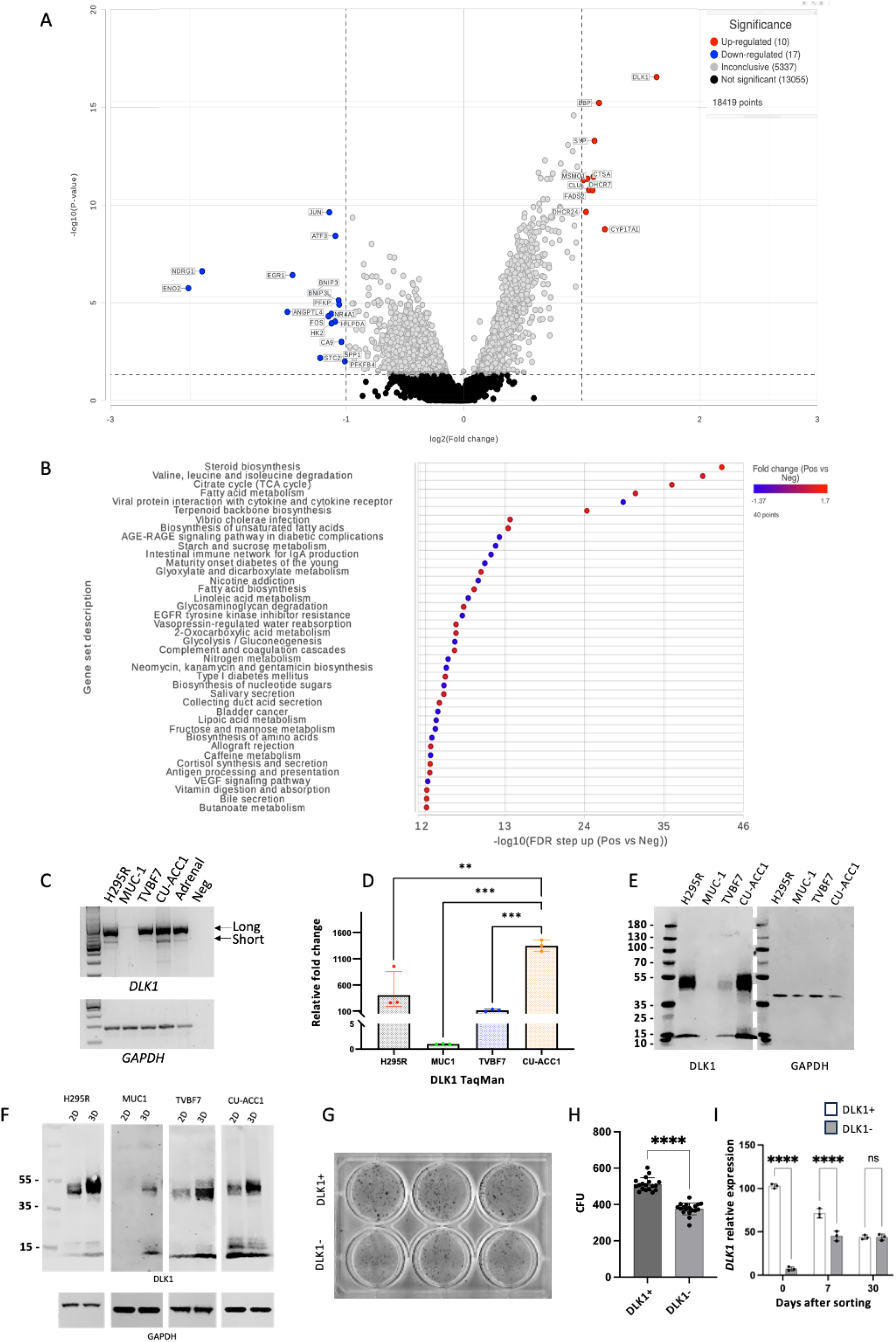
DLK1^+^ cells are endowed with both enhanced steroidogenesis and clonogenicity. A) Volcano plot detailing the genes with altered expression between DLK1^+^ and DLK1^-^ areas of ACC tumors (adj. p <0.01, fold change >/< 2). B) Geneset ANOVA showing the most differentially regulated pathways in DLK1^+^ and DLK1^-^ tumor areas. The most upregulated pathway was of steroid C) PCR analysis of *DLK1* isoforms’ expression in H295R, MUC-1, TVBF7, CU-ACC1 and human adrenal. D) TaqMan analysis of *DLK1* mRNA expression in the indicated ACC lines. E) Western blotting analysis of DLK1 protein expression in the indicated ACC lines. F) Western blotting analysis of DLK1 protein expression in the indicated ACC lines grown in 2D and 3D (spheroids). G-I) Example of colony forming units (CFU) in DLK1^+^ and DLK1^-^ FAC-sorted H295R cells (G) and analysis of CFU (H). I) TaqMan analysis of *DLK1* mRNA expression in DLK1^+^ and DLK1^-^ FAC-sorted H295R cells after sorting, and after 7 and 30 days. **p<0.01, ***p<0.001, ****p<0.0001.

Gene set enrichment analysis and gene set ANOVA were performed. Steroid biosynthesis was the gene ontology (GO) pathway most enriched in the DLK1^+^ group consistent with the upregulation of cholesterol synthesis genes noted above (Figure 6 D-F). There was also enrichment in genes contributing to valine, leucine and isoleucine degradation, TCA cycle and fatty acid (FA) metabolism. Interestingly, apart from CYP17A1, there was no significant differential expression in genes of functional steroidogenesis. Overall, these data support the evidence that DLK1^+^ areas have higher steroidogenic potential compared to Dlk1^-^ areas, primarily through increased cholesterol synthesis, TCA cycle, FA metabolism, as well as direct regulation of steroidogenesis.

Three-dimensional (3D) spheroid cultures are a widely accepted model to enrich cells with (cancer) stem/progenitor features and were used to further assess this apparent paradox of enhanced steroidogenic potential in ACC cells expressing an adrenocortical stem cell marker. We employed four different ACC lines: H295R, MUC-1, TVBF7 and CU-ACC1. All but MUC-1 expressed *DLK1*, and similar to the human adrenal and ACCs, showed a preponderant expression of the full-length *DLK1* (Figure 6 C and D), and detectable DLK1 at the protein level (Figure 6 E). Spheroids could be derived from all cell lines within 14 days in culture (Figure S10 A). DLK1 dosage was significantly enhanced in H295R, TVBF7 and CU-ACC1, and interestingly, *de novo* expression of DLK1 protein was observed in MUC-1 (Figure 6 F). In parallel, mouse BCH-ACC3A cells were also used to generate spheroids (Figure S10 B). LC-MS/MS was then used to compare steroid output of these human and mouse cells in 2D and 3D. 3D spheroids cells showed significant increased output of steroids in H295R, CU-ACC1, BCH-ACC3A and a trend in MUC-1 and TVBF7 (Table S4). To further investigate the co-existence of “stem” (DLK1 expression) and “functional” (steroidogenesis) properties, DLK1^+^ and DLK1^-^ populations were FAC-sorted from H295R cells. Colony forming units (CFU) were assessed after 21 days in culture and DLK1^+^ cells generated significantly more colonies compared with DLK1^-^ cells (Figure 6 G). It is worth noting that expression of *DLK1*, despite being significantly different after sorting, was indistinguishable after 30 days in culture between the two sorted populations (Figure 6 H), suggesting that a steady state of DLK1 expression in cell lines is a prerequisite for viability. These results suggest that ACC cells that express a *bona fide* adrenocortical stem cell marker are endowed with superior steroidogenic potential whilst maintaining some progenitor cell features.

## Discussion

We have previously shown that in rat adrenals, Dlk1 is expressed in Sf1^+^ subcapsular cells and co-expressed with Shh (10) while human adrenals remodel with age to generate DLK1-cell clusters (DCCs). DCCs can be considered “incompletely differentiated” given their phenotype (SF1^+^/CYP11A1^+^/CUP11B2^-^/ CYP11B1^low^/CYP17A1^low^) (11). Here, we showed that in mice, Dlk1 is widely expressed in the adrenal during development. However, cortical expression is restricted to subcapsular clusters as development proceeds, with no cortical expression by P0, while its expression is maintained in the capsule postnatally. Remarkably, Dlk1 is highly expressed in the medulla (as in rats) and currently its role in normal physiology is not known. Capsular Dlk1 expression in postnatal adrenals only partially overlapped with that of PDGFRα (a marker of mesenchymal stem cells), and, although reliable antibodies to Gli1 are not available, widespread expression of both *Gli1* mRNA (RNAScope) and Dlk1 protein suggests the existence of capsular cells which are positive for both markers. Genetic fate mapping has shown Dlk1^+^ cells to be active adrenocortical stem cells during development but near dormant postnatally. Remodeling of the ZG and ZF using dietary restrictions and dexamethasone treatments, respectively, did not result in the generation of Dlk1 progeny postnatally, suggesting a functional diversity of Gli1^+^ and Dlk1^+^ capsular cells.

As i) Dlk1 is capsular postnatally, ii) subcapsular hyperplasia (SH) has capsular-like histological features, iii) SH represents pre-tumoral lesions in some animal models of adrenocortical tumorigenesis and iv) Dlk1 is upregulated in ACC, we hypothesized that SH would be either enriched or derived from Dlk1 expressing cells. However, Dlk1 was not expressed in SH and subsequent tumors in gonadectomized DBA/2J and Inhα/Tag mice. Additionally, SH foci in aged *Dlk1Cre* were derived not from Dlk1 cells. SH has been shown to be derived from Gli1 cells in GDX B6D2F2 mice (37), and indeed in adrenals of aged mice, SH expressed *Gli1* (Figure S4), again suggesting that Dlk1 and Gli1 capsular cells might be, in part, distinct cell populations, at least postnatally. Of note, SH (Sf1^-^/Wt1^+^/Gata4^+^) foci observed in histone methyltransferase *Ezh2* KO mice were shown to be largely derived from the steroidogenic (Sf1^+^) lineage (38).

In an autochthonous ACC mouse model (*BPCre*) combining two major mutations found in patients with aggressive ACC and which closely recapitulates the human disease (15, 16), we have shown DLK1 to be overexpressed in a pattern very similar to human ACC. In *BPCre* mice, targeted expression of activated *Ctnnb1* (β-catenin) and mutated *Trp53* (p53) loss is driven by the AS/Cyp11b2 promoter. Since ZG cells do not express *Dlk1*, *Dlk1* expression is likely re-activated during the development of malignancy but a direct involvement of capsular Dlk1 cells as tumor initiating cells should be ruled out. Re-expression of Dlk1 could therefore represent a “reversion to a stem-like” phenotype which only occurs in sufficiently transformed tissue of a cancer rather than in the renewal and repopulation of functional zones which may occur in homeostasis. It is interesting to note that in spatial transcriptomic analysis, DLK1^+^ tumor areas were programmed to be functional (steroidogenic potential), a finding that was corroborated in human and murine ACC cell lines. When grown in 3D, a recognized experimental set up that leads to enrichment in stem-progenitor cells, there was a significant increase in DLK1 dosage. As recently described, aberrant epigenetic programming in ACC was found to stabilize WNT-active cells which were indeed differentiated (with steroidogenic potential) rather than dedifferentiated as in other cancers (16). Likewise, in this study, the re-expression of a *bona fide* adrenocortical stem/progenitor cell marker, was associated with tumor hormonal “functionality”, further describing the apparent paradox of ACC where differentiation is positively associated with aggressiveness.

In this first prospective study of DLK1 expression in adrenal tumors, there was an incremental level of DLK1 expression, non-adrenal < normal adrenal < adrenal adenoma < ACC. In a further large validation cohort, analyzed retrospectively at a different center, DLK1 expression was found to be a ubiquitous feature of ACC. DLK1 expression was not correlated with other prognostic features, such as Ki-67%, hormonal tumor secretion or ENSAT tumor stage. However higher DLK1 expression was associated with increased risk of disease recurrence. This was most marked in patients with ENSAT stage I-II disease but was also seen in the entire cohort when analyzed by uni- and multivariate analysis controlling for established prognostic factors. The overall effect of DLK1 expression levels on influencing disease progression is more subtle than some of the established markers of worse prognosis (ENSAT stage IV and incomplete resection margins). However, in less advanced disease (ENSAT stages I-III), where complete surgical excision is carried out, DLK1 levels were the strongest factor influencing disease recurrence. This suggests that the metastatic potential of the tumor may be linked to DLK1 expression levels. This would be in keeping with the data from *BPCre* mice where the DLK1 levels were higher in more malignant disease and highest in the metastases. From a functional perspective, this may be partly related to the steroidogenic potential (albeit not necessarily translated to an overt steroid production) of DLK1 positive tumor areas as seen in the spatial transcriptomic analysis. It is known that steroidogenic disease carries worse prognosis in ACC (39, 40). Although DLK1 levels do not differ between groups of disease functionality (Figure 4 F), it is possible that a combination of steroidogenic potential, encompassing tissue level but not clinically detectable steroid hormone excess, and a preponderance of a stem phenotype, classically associated with increase resistance to common oncological treatments are responsible for this, although this needs further investigation.

Additionally, DLK1 levels are measurable in patients with adrenal lesions. Serum DLK1 levels were significantly raised in ACC compared with other benign adrenal adenomas. This was demonstrated to the extent that serum DLK1 levels can predict the diagnosis of ACC in this cohort with a good degree of sensitivity and specificity (serum DLK1 >15.44ng/mL sensitivity 79%, specificity 77%). These findings were validated in a separate cohort from another center, where DLK1 levels were correlated to the disease burden, highest in ENSAT stage IV with the primary tumor *in situ*. This mirrors the findings seen in *BPCre* mice both in tissue and serum. In patients from both centers, where measured, DLK1 levels dropped post adrenalectomy, suggesting that in addition to a possible role in differential diagnosis of adrenal tumors, serum DLK1 levels might be used to longitudinally monitor disease recurrences in patients with ACC. This may be particularly useful in patients who do not have hormonally active disease for whom there are no reliable blood biomarkers for detection of disease recurrence, or those in whom hormone assessments may be complicated by mitotane therapy, and may be adjunctive to the standard radiological surveillance. Further prospective studies are required to confirm these preliminary data.

Despite this consistent finding of raised DLK1 serum levels in ACC, a disparity in mean levels of serum DLK1 was observed between the German and UK centers. This highlights the necessity for further investigation into inter-assay precision. Finally, the positive correlation between DLK1 serum levels and IHC tissue expression, considering the prognostic implication of higher DLK1 levels in tissue, opens the door to further study of DLK1 serum levels in prognosticating disease prior to surgery. Further multicenter prospective studies are required to explore these possibilities but the availability of these measurements of patient serum with a bench top ELISA is enticing as a much more accessible biomarker option than others being proposed in the field.

These data posit Dlk1 positive cells as a novel adrenocortical stem/progenitor marker with an important role in adrenocortical organogenesis and development of malignancy. Expression data in mice and human ACC have shown DLK1 is a marker of increased malignancy and tumor aggressiveness. Furthermore, DLK1 has promise as a biomarker to be used in the diagnosis, prognosis and follow up of patients with ACC. Further larger prospective studies are required to establish this role as are studies looking at DLK1 as a potential therapeutic target in ACC given its preferential expression in this malignancy.

## Methods

| RNAScope reagents and probes | Source | Identifier |
| --- | --- | --- |
| RNAscope® 2.5 High Definition (HD)- RED Assay | ACD | 322350 |
| RNAscope® Probe - Hs-DLK1 - Homo sapiens delta like non-canonical Notch ligand 1 (DLK1) transcript variant 1 mRNA | ACD | 529961 |
| RNAscope® HD Duplex Reagent Kit | ACD | 322430 |
| RNAscope® Probe - Mm-Gli1 - Mus musculus GLI-Kruppel family member GLI1 (Gli1), mRNA | ACD | 31 1001 |
| RNAscope® Probe - Mm-Dlk1-C2 - Mus musculus delta-like 1 homolog (Drosophila) (Dlk1) transcript 1 variant 2 mRNA | ACD | 405971-C2 |

| TaqMan Probes (FAM) | Source | Identifier |
| --- | --- | --- |
| *GAPDH* human | Thermo Fisher | Hs99999905 |
| *DLK1* human | Thermo Fisher | Hs00171584 |
| *Gapdh* mouse | Thermo Fisher | Mm246915_g1 |
| *Dlk1* mouse | Thermo Fisher | Mm00494477_m1 |

| Primary antibodies (IHC and IF) | Host | Source | Identifier | Dilution | AUM |
| --- | --- | --- | --- | --- | --- |
| CYP11B2 (AS 2084) | Rabbit | Celso Gomez-Sanchez, University of Mississippi, USA | N/A | 1:100 | Yes |
| DLK1 (B7) | Mouse | Santa Cruz | Sc-376755 | 1:100 | Yes |
| RFP | Rabbit | Antibodies-online | ABIN129578 | 1:100 | Yes |
| RFP | Goat | Sicgen | AB8181-200 | 1:200 | Yes |
| SF1/NR5A1 (N1665) | Mouse | Invitrogen | 434200 | 1:100 | Yes |
| GATA4 (C-20) | Rabbit | Santa Cruz | sc-1237 | 1:150 | No |
| Tyrosine Hydroxylase | Rabbit | Millipore | AB152 | 1:100 | Yes |
| Ki67 | Rabbit | Abcam | Ab15580 | 1:200 | Yes |
| GFP | Chicken | Abcam | Ab13970 | 1:400 | Yes |
| Active β-catenin | Rabbit | Cell Signaling | 4270S | 1:100 | Yes |
| Cleaved caspase 3 | Rabbit | Abcam | Ab2302 | 1:100 | Yes |

| Secondary Antibodies (IHC) | Host | Source | Identifier | Dilution |
| --- | --- | --- | --- | --- |
| Biotinylated anti Mouse IgG | Goat | Vector | BA-9200 | 1:500 |
| Biotinylated anti Goat IgG | Horse | Vector | BA-9500 | 1:500 |
| Biotinylated anti Rabbit IgG | Goat | Vector | BA-1000 | 1:500 |

| Secondary Antibodies (IF) | Host | Source | Identifier | Dilution |
| --- | --- | --- | --- | --- |
| Alexa Fluor 488 anti chicken IgG | Goat | Invitrogen | A11039 | 1:1000 |
| Alexa Fluor 568 anti mouse IgG | Goat | Invitrogen | A11004 | 1:1000 |
| Alexa Fluor 488 anti rabbit IgG | Goat | Invitrogen | A11008 | 1:1000 |
| Alexa Fluor 568 anti rabbit IgG | Goat | Invitrogen | A11036 | 1:1000 |
| Alexa Fluor 488 anti mouse IgG | Goat | Invitrogen | A11029 | 1:1000 |

| Primary antibodies<br>(Western Blotting) | Host | Source | Identifier | Dilution |
| --- | --- | --- | --- | --- |
| DLK1 (N18) | Goat | Santa Cruz | sc-8623 | 1:500 |
| DLK1 (B7) | Mouse | Santa Cruz | sc-376755 | 1:100 |
| HA (Y11) | Rabbit | Santa Cruz | sc-805 | 1:500 |
| FLAG (M2) | Mouse | Merck | F1804 | 1:500 |
| GAPDH | Mouse | Santa Cruz | sc-47724 | 1:2000 |

| Secondary antibodies<br>(Western Blotting) | Host | Source | Identifier | Dilution |
| --- | --- | --- | --- | --- |
| IRDye 800 CW anti rabbit IgG | Goat | Li-cor | 926-32211 | 1:10000 |
| IRDye 680 RD anti mouse IgG | Goat | Li-cor | 926-68070 | 1:10000 |
| IRDye 680 RD anti goat IgG | Donkey | Li-cor | 925-68024 | 1:10000 |

### Genetic lineage tracing

Mice were housed in a 12 h light/12 h dark cycle in a temperature- and humidity-controlled room (21 °C, 55% humidity) with constant access to food and water. Experimental procedures in the UK were under the terms of a UK government Home Office license (PPL P48019841). All mice were maintained on a C57BL/6 background and included Rosa26^CAGLoxpSTOPLoxpTdTomato^ (RRID:IMSR_JAX:007914), Dlk1CreERT2 (a gift from Prof Fiona Watt, Kings College, London, UK), PDGFRα-H2BEGFP (RRID:IMSR_JAX:007669). Rosa26^CAGLoxpSTOPLoxpTdTomato^ mice were crossed with Dlk1CreERT2 mice to generate DLK1-CreER, Rosa26C^TdTomato/+^ mice. Axin2Cre:ERT2/+ mice and RosaYFP/YFP mice were purchased from Jackson laboratories. These mice were crossed to produce Axin2CreERT2/+; RosaYFP/YFP mice for lineage tracing studies. Tamoxifen (200mg/g in corn oil, given via IP or orally) was given to dams or postnatal mice, and chase times varied as described in the main text. Initial experiments were aimed at assessing Tamoxifen dose resulting in >80% recombination after 6 days, assessed by immunohistochemistry (IHC) on consecutive sections with anti-RFP and anti-Dlk1. No leakage was observed in random adrenals taken from sham injected Dlk1CreERT2, Rosa26C^TdTomato/+^ mice, and stained with anti-RFP.

For ZF remodeling, two inductions regiments were used: 1. p60 (to 7 months) and p460 *Dlk1Cre* mice were treated with tamoxifen (200mg/g in corn oil, oral gavage) on day 0 and day 3 and with dexamethasone (6.5 μg/g in corn oil, oral gavage) or vehicle on day 1,2,4 and 5. 2. p30 and p460 *Dlk1Cre* mice were treated with tamoxifen on day 0, day 3 and day 6, and with dexamethasone or vehicle on day 1,2,4 and 5. Corticosterone was measured at day 0, 6 and 21 (Enzo Life Sciences).

For ZG remodeling, p50/70 *Dlk1Cre* mice were assigned to experimental groups according to the different sodium chloride contents in the chow (standard diet, low sodium 0.003%, high sodium 3.3%, SAFE^®^ Complete Care Competence) for 8 days, with tamoxifen injection (200mg/g in corn oil, oral gavage) at day 0 and at day 3.

Adult mice underwent transcardiac perfusion with phosphate-buffered saline (PBS), followed by paraformaldehyde (PFA). Embryos/postnatal adrenals were fixed/postfixed in PFA, before paraffin embedding.

Timed pregnancies were achieved by mating females and males overnight and, the presence of vaginal plug the following morning, was considered as embryonic day (e) 0.5.

### Murine ACC model and ACC cell line

The protocols of animal experiments were approved by Boston Children’s Hospital’s Institutional Animal Care and Use Committee. The *BPCre* mouse model of spontaneous ACC was bred as described previously(15). These mice express activated *Ctnnb1* (β-catenin) and mutated *Trp53* (p53) (*AS^Cre/+^*:: *Trp53^flox/flox^*:: *Ctnnb^flox(ex3)/+^*) in the adrenal and spontaneously develop metastatic ACC. Additionally, the derivation of the BCH-ACC3A cell line from a *BPCre* tumor is described elsewhere (16). Tumors were weighted, before being fixed in PFA and embedded in paraffin.

Following retro-orbital blood collection, serum was stored at -80°C until analysis. Samples were thawed and analyzed using the Mouse Dlk1 ELISA Kit (Invitrogen EM66RB) following the manufacturer’s instructions.

### Subcutaneous tumor model

This study was performed in compliance with Home Office PPL PP6127261. H295R in exponential grow were collected and cell suspensions (10 × 10^6^ cells/100 μl in 10% Tween-80 PBS) were inoculated subcutaneously into the right flanks of 9-week-old female NMRI-Foxn1nu/nu mice (Janvier labs). Tumor take rate was 80%. Tumor volume (mm^3^) was assessed via caliper measurement twice a week and was calculated by using the formula: length x width^2^/2. Pentobarbital anesthetized mice were exsanguinated by cardiac puncture when tumors reached different sizes, blood was collected for serum human DLK1 measurements (AdipoGen life Sciences). Tumors were also measured after collection before being fixed in PFA, embedded in paraffin and sections processed for DLK1 IHC.

### Gonadectomized mice model

The University of Turku Ethical Committee on Use and Care of Animals approved all procedures of the current experiments. Maintenance of DBA/2J and inhibin α subunit promoter (Inhα)/Simian virus 40 T-antigen mice and gonadectomy procedures were described previously (26, 27).

### Human Tissues collection and processing

Human adrenal specimens were collected from patients undergoing surgery at each of St Bartholomew’s, University College and Hammersmith Hospitals, London, after written consent obtained from participants and under the study protocol *Genetics of endocrine tumors* (REC: 06/Q0104/133).

In Germany, all tissue was collected under the ENS@T research ethical agreement (No. 88/11) at the Universitätsklinikum Würzburg. All patients gave informed consent. All clinical data were collected through the ENS@T database (registry.ensat.org).

Samples were fixed in 4% paraformaldehyde (PFA) for 10-24 hours at 4°C and embedded into paraffin. Sections were cut at 2-8μm using a rotary microtome (Thermo scientific) and transferred onto SuperFrost Plus Adhesion slides (VWR).

### Immunohistochemistry and section analysis

Formalin-fixed paraffin-embedded (FFPE) sections were deparaffinized in xylene (3 x 10-minute washes), washed in 100% ethanol (2 x 10-minute washes) and incubated in 3% hydrogen peroxide solution in methanol for 30 minutes at room temperature (RT) to block endogenous peroxidase activity. After dehydration in a descending ethanol series (100, 90, 70, and 50% each concentration for 10 minutes), sections were washed in ddH*2*O, submerged in citrate buffer (Vector) for 20 mins at 95°C and then allowed to gradually reach RT. They were then blocked with 10% goat serum in PBS-Triton 0.1% (T-PBS) containing 4 drops/ml of Avidin solution (of the Avidin/Biotin Blocking Kit, Vector Labs SP-2001) for 1 hour and then incubated overnight with the primary antibody (Table 1) containing 4 drops/ml of Biotin Solution (of the Avidin/Biotin Blocking Kit, Vector Labs) at RT. Slides were washed with T-PBS and incubated with biotinylated goat anti-rabbit secondary antibody (Table 1) diluted in T-PBS for 2 hours at RT. After further washes in T-PBS slides were incubated with the Avidin-Biotin Complex (Vector Labs, PK-6100) at RT for 1 hour. Following washes sections were developed with 3,3’-diaminobenzidine (Vector Labs) and counter-stained with Gill hematoxylin (Sigma). Slides were dehydrated, incubated with xylene and mounted using Vectamount mounting medium (Vector Labs).

In Germany, IHC was performed on full sections of each tumor sample. Slides were deparaffinized in xylene (2 x 10-minute washes) and rehydrated in ethanol (100, 90, 80, and 70% each concentration for 5 min.). Slides were washed 5 times in ddH2O before high temperature antigen retrieval was performed in 10 mM citric acid monohydrate buffer (pH 6.5) (Sigma) in a pressure cooker (Silit) for 13 min. After 20 minutes cooling at RT, slides were washed five times in ddH2O, and endogenous peroxidase activity was blocked for 10 minutes in the dark with 3% hydrogen peroxide solution in methanol. After five washes in ddH2O, blocking of unspecific protein-antibody interactions was performed with 20% human AB serum (Sigma) in PBS for one hour at RT in the dark. Primary antibody was then added in PBS and slides were incubated at RT for one hour. After five washes in PBS, signal amplification was achieved by HiDef DetectionTM HRP Polymer System for 20 minutes at RT as per manufacturer’s instructions. After 3 x 2 minute washes in PBS, slides were then developed for 10 min with DAB Substrate Kit (Vector Labs) according to the manufacturer’s instructions. Development was stopped by three washes in tap water. Nuclei were counterstained with Mayer’s hematoxylin for three minutes. Slides were then washed for 2 minutes in running tap water before dehydration for 2 minutes sequentially in 70%, 100%, 100% ethanol and xylene. Finally, slides were mounted with Entellan (Merck).

In London, slides were scanned at 20x magnification with a Grundium Ocus slide scanner (Grundium). Scanned images were imported into Qupath (Open-source software for digital pathology image analysis (41) and manually annotated. Positive cell detection software was run to generate a H score /300 for each section.

In Germany, slides were scanned at 20x magnification on an Aperio Versa microscope (Leica Biosystems, Germany). Images were checked, manually annotated, and then analyzed using Aperio Positive Pixel Count software (Leica). Staining intensity and distribution were calculated by the software and a H score /300 was generated for each sample.

### Immunofluorescence

For immunofluorescence, the protocol for IHC minus the hydrogen peroxide step and up to the blocking step was followed. Sections were incubated with primary antibodies overnight at RT, washed in T-PBS, incubated with fluorescently labelled secondary antibodies and reacted with 4′,6-diamidino-2-phenylindole (DAPI, Sigma) before mounting. Images were acquired using a Leica DM5500B microscope, equipped with a DCF365FX camera (Leica), and then processed with Abode Photoshop CS6.

### Cell culture maintenance

HEK-293T cells were maintained in DMEM high glucose (Gibco) and 10% fetal bovine serum (FBS).

NCI-H295R cells were maintained in DMEM/F-12 HAM (1:1)/GlutaMAX (Gibco), 1% Insulin-Transferrin-Selenium (Scientific lab) and 2.5% NuSerum (Scientific lab).

CU-ACC1 cells were grown in F12 Nutrient ham (Gibco) and DMEM-high glucose, pyruvate (Gibco) at 3:1 V/V ratio containing 10% FBS, 0.4ug/ml Hydrocortisone, 5ug/ml Insulin, 8.4ng/ml Cholera toxin, 24ug/ml Adenine, 10ng/ml EGF. TVBF7 cells were grown in DMEM/F-12 HAM (1:1) + GlutaMAX (Gibco) and 10% FBS.

MUC1 cells were maintained in DMEM/F-12 HAM (1:1) + GlutaMAX (Gibco) and 10% FBS. All cell lines were supplemented with 1% Penicillin-streptomycin and cultured in 5% CO_2_ at 37°C. Cells were confirmed to be mycoplasma free by monthly testing using the MycoAlert Detection Kit (Lonza).

### FAC-sorting

H295R and HEK-293T cells were dissociated with trypsin-EDTA and suspended in 20ml of new complete medium in T75 cell suspension flasks (Cellstar) overnight. The following day cells collected by centrifugation at 1000 x g for 5 minutes, resuspended in 5ml medium, and passed through a 40μm cell strainer before counting with a hemocytometer. Samples were separated into 1.5ml Eppendorf tubes (1 x unstained sample, 1 x DAPI, 1 x sorting sample). At least 50,000 cells were used for the unstained and DAPI and the rest saved for sorting. Tubes were taken to the bench and spun at 1000 x g for 5 minutes. Supernatant was aspirated and cells were resuspended in new tubes in 0.5ml sterile FACS buffer (50ml PBS, 0.5g bovine serum albumin (BSA), 2mM EDTA). Tubes were centrifuged in same conditions as above and supernatant was discarded. Cells were resuspended in 200μl of FACS buffer and conjugated antibody was added to the sample for sorting (Human Pref-1/DLK1/FA1 Alexa Fluor® 488-conjugated Antibody (R&D Systems)) at recommended concentration of 5μL/10^6^ cells. At the same time, 0.5μL of antibody was added to UltraComp eBeads™ Compensation Beads (Thermo Fisher) in 200μL of FACS buffer. Samples were incubated on ice in the dark for 30 minutes, being vortexed every 10 minutes). All samples were washed 3 times with 1ml FACS buffer and spun at 1000 x g for 5 minutes. DAPI was added to the single DAPI control and the sample for sorting at a dilution of a 0.1mg/ml solution. All samples were passed through another 40μm cell strainer into polystyrene FACS tubes (Corning) and taken to the Flow Cytometry Facility in William Harvey Research Institute, QMUL. Staff in the facility optimized the settings and carried out the sort as per departmental protocol using a BD FACSAria II. Gating was initially optimized using non transfected and DLK1 transfected HEK293T cells. Sorted samples were collected in polystyrene FACS tubes (Corning) containing 0.5ml FACS buffer. DLK1^+^ and DLK1^-^ FAC-sorted H295R cells were immediately plated in 6 well-plates at a density of 3×10^3^ cells/well and cultured for 3 weeks, after which the number of colonies in each plate was counted manually. Sorted cells were also processed for RNA extraction.

### Spheroid generation

Human H295R, CU-ACC1, MUC-1, TVBF7, and murine BCH-ACC3A cells were plated at 4- 5×10^3^ cells per well in ultra-low attachment 6 well plates (Corning) in spheroid medium (DMEM/Nutrient Mixture F-12 Ham (Sigma) supplemented with recombinant human basic fibroblast growth factor (20 ng/mL) (Sigma), recombinant human epidermal growth factor (20 ng/mL) (Sigma), B-27 (Thermo Fisher), N-2 supplements (Thermo Fisher)), and spheroids were allowed to form over a period of 14 (H295R, CU-ACC1, MUC-1 and TVBF7) or 7 days (BCH-ACC3A). Bright field pictures were taken with am AxiovertA1 microscope (Zeiss).

Medium was collected, centrifuged for steroid analysis, using total RNA extracted from spheroids and 2D cultures for normalization. In parallel experiments, spheroids were allowed to fall by gravity in 15 ml tubes, washed with PBS, and then lysed with RIPA buffer prior to western blotting.

### RNAScope

Sections were processed to detect human *DLK1* mRNA using the RNAScope 2.5 High Definition (HD)-RED Assay (Advanced Cell Diagnostics) and for mouse *Dlk1* (Red) and *Gli1* (Green) mRNAs using the RNAscope HD Duplex Reagent Kit according to the manufacturer’s instructions.

### Cloning of DLK1 isoforms

Primers spanning the human *DLK1* cDNA sequence FW 5’- <u>cggaattcag</u>ATGACCGCGACCGAAGCC-3’ (<u>EcoRI</u>) and REV 5’- <u>ccgctcgag</u>TTAGATCTCCTCGTCGCC-3’ (<u>XhoI</u>) were used for PCR on cDNA from human adrenal (n=3, pulled) and H295R cells, using Platinum™ *Taq* DNA Polymerase, high fidelity (Thermo, denaturation 94^0^C 30 seconds, 35 cycles 94^0^C 15 seconds, 60^0^C 30 seconds, 68 ^0^C 2 minutes, final extension 68 ^0^C 5 minutes). The two amplicons (924bp and 1149bp) were subcloned into pCMV-HA vector (Clontech, K6003-1) and sequenced, resulting in the cloning of both the full length and shorter DLK1 isoforms. Using the resulting vectors as templates, both isoforms were subcloned into pCMV-Tag4 (Stratagene, 211174) using primers FW <u>cgcggatcc</u>ACCATGTACCCATACGATG (<u>BamHI</u>) and REV <u>ccgctcgag</u>GATCTCCTCGTCGCCGGC (<u>XhoI</u>) (PCR conditions as above). The final vectors encoded HA (N-term) and FLAG (C-term)-tagged DLK1 proteins.

### *DLK1* isoforms PCR

Primers to simultaneously detect the full length and shorter human *DLK1* isoforms were FW 5’-AACAACAGGACCTGCGTGAG-3’ and REV 5’-GCAGGTTCTTCTTCTTCCGCA-3’, with amplicon sizes of 754bp and 535bp, respectively. New England Biolabs Hot Start *Taq* DNA Polymerase was used, and PCR cycle was as follows: denaturation 95^0^C 30 seconds, 35 cycles 95^0^C 20 seconds, 60^0^C 30 seconds, 68 ^0^C 30 seconds, final extension 68 ^0^C 5 minutes.

### ELISA of patient serum/plasma

In London, blood was taken from patients pre-operatively or at the start of chemotherapy treatment in the neo-adjuvant or non-operative management setting. Blood was taken post-operatively at the first outpatient appointment. All blood draws included a yellow SST bottle for serum. This was allowed to clot at RT for 10-15 minutes before being spun at 4°C at 1000 x g for 10 minutes and stored in 200-500μl aliquots at -20°C. When possible, blood draws also included a purple EDTA bottle for plasma. These samples were spun and stored as above. In Germany, samples were processed and stored as per local protocols and guidelines. Samples were identified for analysis and aliquoted in 200μl volumes for analysis. Serum and plasma samples were run and analyzed with the DLK1, Soluble (human) ELISA Kit (Adipogen) as per manufacturer’s instructions.

### Transfection

Human embryonic kidney cells (HEK293-T) cells were transfected using Lipofectamine™ 3000 Transfection Reagent (Invitrogen) accordingly to the manufacturer’s instructions. Forty-eight hours post transfection, cells were lysed, protein extracted, quantified and used for western blotting. In some experiments, medium was changed to serum-free medium 24 hours post transfection and collected 72 hours post transfection for DLK1 ELISA (Adipogen).

### Protein extraction

Samples were lysed in cold RIPA lysis buffer (Thermo Fisher) supplemented with protease inhibitor cocktail (Roche). Cells were firstly washed in PBS, and directly lysed in buffer, whilst freshly isolated adrenals were minced using a Precellys 24 homogenizer (Bertin Instruments) with Precellys Lysis kit in lysis buffer. Lysates were kept on ice for 20 minutes and then cleared by centrifugation at 4C for 10 mins at 13,000 RPM. Protein concentration was determined using the BCA kit (Pierce).

### Western blotting

Protein samples (20 μg) were size-separated on 4-12% NUPAGE gels (Thermo Fisher), and gels blotted onto nitrocellulose membranes (Protran). Membranes were stained with Ponceau to assess equal loading, de-stained in PBS containing 0.1% Tween-20 (PBS-T), blocked with 5% non-fat dry milk in PBS-T and incubated with primary antibody overnight at 4°C. After washes in PBS-T, membranes were incubated with secondary antibodies. Immunoblots were scanned using the Odyssey XF Imaging System (LI-COR).

### RNA extraction and cDNA synthesis

Total RNA was extracted using the RNeasy® Mini kit (Qiagen) according to the manufacturer’s instruction. During extraction DNA was digested with DNaseI for 15 minutes at RT (Qiagen). RNA concentration and RNA quality (A260/A280 ratio) was determined using a nanodrop (Thermo Fisher). 500ng of RNA in a 20 ul reaction was reverse transcribed into cDNA using a High-Capacity cDNA Reverse Transcription (Applied Biosystems), according to the manufacturer’s instruction.

### Real Time qPCR

Real-time quantitative PCR was performed using TaqMan® Universal Master Mix II and TaqMan® assays (Applied Biosystems, ABI). Pre-made primers and FAM probes were purchased from Thermo Fisher. The final reaction volume of 20μL consisted of 10μL TaqMan® Universal Master Mix II (2X), 1μL TaqMan® Gene Expression Assay (2X) and 9μL 2.5ng/mL cDNA template. Amplification and detection were performed with the AriaMx Real-time PCR System with the following cycle condition: 95°C for 10 min, 40 cycles at 95 °C for 10 sec and 60 °C for 1 min. Each measurement was carried out in triplicate. Differences in gene expression, expressed as fold-change, were calculated using the 2−ΔΔCt method using *Gapdh* as the internal control.

### GeoMx spatial transcriptomics

Complete methods for GeoMx assays can be found in (6) and in the GeoMx manual. Four FFPE ACC samples were used for spatial transcriptomics. For each block, serial 5 μm sections were mounted on to superfrost plus slides and the best consecutive five processed as follow: slide 1 for RNA quality control after scraping the sections off, slide 2 for H&E, slide 3 for DLK1 IHC, slide 4 was the experimental one, slide 5 for *DLK1* RNAScope. The experimental slides were backed in a 60°C drying oven for 1 hour, deparaffinized and antigen unmasking was performed in citrate buffer pH 6.0 in a pressure cooker. Slides were then allowed to cool. The mix of Whole Transcriptome Atlas probes (WTA, Nanostring) was dropped on each section and covered with HybriSlip Hybridization Covers. Slides were then incubated for hybridization overnight at 37°C in a Hyb EZ II hybridization oven (Advanced cell Diagnostics). The day after, HybriSlip covers were gently removed and 25 minutes stringent washes were performed twice in 50% formamide and 2X saline sodium citrate (SSC) at 37 °C. Slides were washed for 5 min in 2XSSC, then blocked in Buffer W (Nanostring) for 30 min at RT in a humidity chamber, washed in T-PBS and blocked with buffer W, before overnight incubation in a humidity chamber at 4°C with anti DLK1 (B7, Santa Cruz Biotechnologies) diluted 1:100 in buffer W. After washes in 2XSSC, sections were incubated with secondary antibodies (Goat anti mouse Alexa Fluor 488, Invitrogen, 1:500 dilution in buffer W) for 1 hour and nuclei were stained with SYTO 13 (Nanostring). Sections were then loaded in a GeoMx DSP instrument, scanned and 60 regions of interest (ROI) chosen, based on DLK1 IF signal. The DLK1^+^ and DLK1^-^ ROI included only tumor cells based on morphological and histological examination of the slide and the adjacent H&E, RNAScope and IHC slides.

ROI were then exposed to 385 nm UV light allowing release of the indexing oligos which were then collected in a 96-well plate. Oligos were dried and resuspended in 10 μL of DEPC-treated water. Sequencing libraries were generated by PCR from the photo-released indexing oligos and AOI-specific Illumina adapter sequences and unique i5 and i7 sample indices were added. Each PCR reaction used 4 μL of indexing oligos, 1 μL of indexing PCR primers, 2 μL of Nanostring 5X PCR Master Mix, and 3 μL PCR-grade water. Thermocycling conditions were 37°C for 30 min, 50°C for 10 min, 95°C for 3 min; 18 cycles of 95°C for 15sec, 65°C for 1min, 68°C for 30 sec; and 68°C 5 min. PCR reactions were pooled and purified twice using AMPure XP beads (Beckman Coulter, A63881) according to manufacturer’s protocol. Pooled libraries were sequenced at 2×75 base pairs and with the single-index workflow on an Illumina NextSeq to generate 458M raw reads. Data was analyzed using the Partek software (Illumina).

### Steroid hormone quantification

Quantification of steroids was achieved by liquid chromatography-tandem mass spectrometry (LC-MS/MS).

#### Calibrator, Internal Quality Control and Internal Standard solutions

Certified reference material (Cerilliant, Merck) stock solutions for all analytes (each 1000 mg/L, except pregnenolone, 11-deoxycorticosterone, cortisone and 21-deoxycortisol, all 100 mg/L) were used. These were used to prepare combined working solutions containing all analytes for calibration and internal quality control (IQC) purposes. To make these, appropriate volumes of each stock solution were added to a glass tube and then dried down under nitrogen at 60°C. The steroids were then reconstituted in methanol to create calibrator and IQC working solutions each containing: DHEAS (4000 μg/mL), cortisol (200 μg/mL), 17-hydroxypregnenolone (40 μg/mL) 17-hydroxyprogesterone (40 μg/mL), androstenedione (12 μg/mL), pregnenolone, corticosterone, 11-deoxycortisol, 21-deoxycortisol, cortisone (each at 40 μg/mL), testosterone (8 μg/mL) and 11-deoxycorticosterone (8 μg/mL). The working solutions were each further diluted in methanol to create three further working solutions as follows: 3+20 (v/v), 1+39 (v/v) and 1:199 (v/v). All four working solutions were then used to make appropriate volumes of calibration standard/IQC solution by dilution in DMEM. After thorough mixing and equilibration (24 h, 2–8 °C), calibrators and IQC solutions were portioned in 1.5 mL microcentrifuge tubes (Eppendorf, Stevenage, UK) and stored at -20 °C until required.

Internal standards (IS) stock solutions were prepared in methanol (each 1000 mg/L). A combined IS sub-stock solution was prepared in methanol containing deuterated steroids at the following concentrations: DHEAS-D2 (75000 μg/mL), cortisol-D4 (15000 μg/mL), 17-hydroxypregnenolone-D3 (1500 μg/mL) 17-hydroxyprogesterone-D8 (500 μg/mL), androstenedione-D7 (150 μg/mL), pregenenolone-D4 (3000 μg/mL), cortisone-D2 and corticosterone-D8 (both at 1500 μg/mL), 11-deoxycortisol-D2 and 21-deoxycortisol-D8 (both at 200 μg/mL), testosterone-D3 (300 μg/mL) and 11-deoxycorticosterone-D8 (50 μg/mL). The IS working solution was freshly prepared before each batch by dilution of 2.5 μL per mL of IS working solution required in methanol.

#### Specimen processing

Portions of frozen calibrators, IQC solutions and unknown media samples were thawed and mixed at RT by inversion before analysis. Aliquots (50 μL) of calibrator /IQC/unknown media samples were transferred into 1.5 mL micro-centrifuge tubes and 100 µL of IS working solution added to each tube. Tubes were capped, vortex-mixed for 5 seconds and then 200 μL of deionized water was added to each tube. Tubes were once again capped and vortex-mixed for 5 seconds and then centrifuged (13,000 g, 5 min). 300 μL of supernatant was then added to individual wells of an Oasis Max µElution solid phase extraction (SPE) plate (Waters Corp). Each SPE well had earlier been preconditioned with 150 μL of methanol, followed by 150 μL of deionized water. Subsequently, each SPE well was washed with 100 µL of 1% (v/v) formic acid in 15% acetonitrile (aq), followed by 100 µL of 1% (v/v) ammonia in 15% acetonitrile (aq). Finally, captured steroids were eluted into a 96 well plate by adding 50 µL of 60% acetonitrile (aq) to each SPE well. 50 µL deionized water was then added to each well of the collection plate.

#### LC-MS/MS procedure

LC-MS/MS was performed using a 1290 Infinity II LC System coupled with a 6495 triple quadrupole mass spectrometer (both Agilent Technologies). Extracts were injected (5 μL) onto the LC column (Zorbax Eclipse Plus C18 2.1×50mm, 1.8 µm) at a flow rate of 0.6 mL/min at 40°C. Mobile phases were (A) 1 mM ammonium fluoride in 60:40 (v/v) dH_2_O: methanol and (B) 1 mM ammonium fluoride in methanol. MS/MS was carried out in positive mode using electrospray ionization (ESI; Gas temp: 230°C, Gas Flow: 16 L/min, Nebulizer: 25 Psi, Sheath Gas Temp: 400°C, Sheath Gas Flow:12 L/min, Capillary: 4500 V, Nozzle voltage: 4500 V) operated in selected reaction monitoring (SRM) mode, with two m/z transitions per analyte and one m/z transition for each internal standard. LC-MS/MS instrument control, data acquisition and post-analysis processing was performed using MassHunter (version B.09.00, Agilent Technologies). For assay calibration, peak area ratios (analyte quantifier to IS) were used to construct calibration graphs, with lines fitted by linear regression. The intercepts were not forced through zero, and line weighting was applied (1/concentration). Deuterated ISs were used for all steroids in the developed method.

### Statistical analysis

All data are presented as mean + standard deviation (SD) unless otherwise stated. Fisher’s exact test or χ2 test was used to investigate dichotomic variables, whereas two-sided student’s t-test and Pearson correlation were used to test continuous variables. Where multiple comparisons were made, a one-way ANOVA was performed followed by post hoc Tukey’s multiple comparison tests to generate adjusted p values. Correlations and 95% CIs between different parameters were evaluated by linear regression analysis. Overall survival (OS) was defined as the time from the date of primary surgery to specific death or last follow-up, whereas recurrence-free survival (RFS) was defined as the time from the date of primary tumor resection, after complete resection (R0), to the first radiological evidence of any kind of disease relapse or death. Progression-free survival (PFS) was defined as the time from the date of data capture to the first radiological evidence of any kind of disease progression or relapse or death. All survival curves were obtained by Kaplan–Meier estimates, and the differences between survival curves were assessed by the log-rank (Mantel–Cox) test. For the calculation of hazard ratios (HRs), two ACC groups with low or high DLK1 expression were considered (higher or lower than median DLK1 expression). Additionally, four ACC groups with low, low-intermediate, high-intermediate, and high DLK1 expression were considered based on DLK1 expression in quartiles. A multivariate regression analysis was performed by Cox proportional hazard regression model to identify factors that might independently influence survival. Statistical analyses were made using GraphPad Prism version 9.1 (La Jolla, CA, USA) and SPSS Software PASW Version 26.0 (SPSS, Inc., Chicago, IL, USA). P values of <0.05 were considered statistically significant.

## Acknowledgments

This work was supported by the MRC (MR/X021017/1 to LG), BBSRC (BB/V007246/1 to LG), Barts Charity (MGU0436 to LG), Rosetrees Trust (M355-F1 to LG), MRC (MR/S022155/1 to JFP), the German Research Foundation (Deutsche Forschungsgemeinschaft (DFG), grant agreement number: 314061271 to MF), R01DK123694 (to DTB).

We would like to thank all the patients, both in UK and in Germany, who consented to the use of their samples and clinical information in this study, especially the family of Joanne Baldock, who donated money to the department for a piece of equipment to further research in ACC.

## Author contributions

Conceptualization LG, JFHP, KM

Methodology LG, JFHP, KM, BA, SSb, MK, MF, WD, KSB, DB

Validation KSB, CR, KM, JFHP, BA, JAL

Formal analysis JFHP, KM, BA, KSB

Investigation OR, DRT, JP, KM, BA, LG, SSb, ER

Resources SSi, AA, MT, KKV, MW, LP, TEAA, TTC, ADM, FP, CGS, CH, JF, JC, JS, NR, MD

Data curation JFHP, KM, LG

Writing – original draft JFHP, KM, LG

Writing – review and editing: all authors.

Supervision LG, WMD, MF, MK, DB

Project administration LG

