## Supplementary Tables and Figures for "Dlk1 is a novel adrenocortical stem/progenitor cell marker that predicts malignancy in adrenocortical carcinoma"

| | | n | Median | Range |
|---|---|---|---|---|
| **Sex** | Male | 26 | | |
| | Female | 47 | | |
| | Total | 73 | | |
| **Age at operation (years)** | | | 52 | (17-77) |
| **Diagnosis** | ACC | 19 | | |
| | APA | 12 | | |
| | NAPACA | 28 | | |
| | Phaeochromocytoma | 1 | | |
| | Other benign | 9 | | |
| | Other malignant | 4 | | |

**Table S1. Descriptive characteristics of London patient cohort**
ACC– adrenocortical carcinoma, APA – aldosterone producing adenoma, NAPACA – non-aldosterone producing adrenocortical adenoma. The “other benign” category includes histological diagnoses of: micronodular hyperplasia, nodular and diffuse hyperplasia, ganglioneuroma, pseudocyst, focal nodular hyperplasia of zona glomerulosa, retroperitoneal schwannoma, perivascular epithelial cell tumor (PEComa), florid hyperplasia with myelolipomatous metaplasia and a sample characterized by fibrotic material with calcification. The “other malignant” category includes liposarcoma, 2 metastases from adenocarcinoma of the lung and a metastasis from ovarian carcinoma.

| **Overall Survival** Variables | **n** | **Median survival (months)** | **Univariate** | | | **Multivariate** | | |
|---|---|---|---|---|---|---|---|---|
| | | | **HR** | **95% CI** | **p** | **HR** | **95% CI** | **p** |
| **Age (years)** | | | | | | | | |
| 0-49 | 72 | 46 | | | | | | |
| 50+ | 58 | 47 | 0.991 | 0.6409 to 1.517 | 0.9673 | | | |
| **ENSAT stage** | | | | | | | | |
| I-II | 56 | 76 | | | | | | |
| III | 44 | 49 | 1.339 | 0.7963 to 2.256 | 0.2697 | 1.293 | 0.7421 to 2.254 | 0.3627 |
| IV | 30 | 20 | 3.452 | 2.011 to 5.919 | **<0.0001****** | 1.537 | 0.7338 to 3.295 | 0.2608 |
| **Ki-67%** | | | | | | | | |
| 0-19 | 58 | 65 | | | | | | |
| 20+ | 65 | 30 | 1.71 | 1.091 to 2.705 | **0.02**** | 1.537 | 0.9526 to 2.503 | 0.0804 |
| **Glucocorticoid excess** | | | | | | | | |
| Absent | 38 | 45 | | | | | | |
| Present | 57 | 39 | 1.398 | 0.8449 to 2.380 | 0.2022 | | | |
| **Resection status** | | | | | | | | |
| 0 & X | 89 | 75 | | | | | | |
| 1 & 2 | 40 | 22 | 3.72 | 2.351 to 5.865 | **<0.0001****** | 3.279 | 1.681 to 6.169 | **0.0003***** |
| **DLK1 expression** | | | | | | | | |
| Low | 65 | 47 | | | | | | |
| High | 65 | 46 | 1.081 | 0.7061 to 1.656 | 0.7188 | 1.086 | 0.6872 to 1.722 | 0.7236 |

**Table S2. Cox regression analysis of overall survival in validation cohort (Würzburg)**

| | | n | Median | Range |
|---|---|---|---|---|
| **Sex** | Male | 8 | | |
| | Female | 17 | | |
| | Total | 25 | | |
| **Age at sample (years)** | | | 51 | (26-80) |
| **Disease status at time of sample** | Primary tumor | 9 | | |
| | Primary tumor and metastases | 6 | | |
| | Recurrence after primary surgery | 17 | | |
| | Disease free | 7 | | |
| | Total | 39 | | |
| **Hormone secretion** | Cortisol (alone or mixed) | 12 | | |
| | Androgens | 2 | | |
| | Inactive | 8 | | |
| | Unknown | 2 | | |
| **ENSAT tumor stage at time of sample** | I | 0 | | |
| | II | 3 | | |
| | III | 3 | | |
| | IV | 22 | | |
| **Tumor size (cm)** | Primary tumor | (15) | 13 | (6.5-19) |
| | Local recurrence | (3) | 4 | (3.5-7) |
| **Resection status** | 0 | 7 | | |
| | 1 | 2 | | |
| | 2 | 5 | | |
| | X | 2 | | |
| | N/A | 3 | | |
| **Weiss score** | | | 8 | (5-10) |
| **Ki-67 %** | | | 20 | (10-70) |

**Table S3. Descriptive characteristics of validation cohort for serum analysis (Würzburg)**

| | | 2D | | | | 3D | | | | | |
|---|---|---|---|---|---|---|---|---|---|---|---|
| **Cell line** | **Steroid** | **Mean normalized concentration (nmol/L/RNA mass) ± SD** | | | **n** | **Mean normalized concentration (nmol/L/RNA mass) ± SD** | | | **n** | **Significance** | **p value** |
| **H295R** | Corticosterone | 0.000 | ± | 0.000 | 3 | 1.485 | ± | 0.4042 | 3 | ** | 0.0031 |
| | 11-deoxycortisol | 6.170 | ± | 1.609 | 3 | 14.960 | ± | 3.890 | 3 | * | 0.0224 |
| | Androstenedione | 4.800 | ± | 1.349 | 3 | 1.934 | ± | 0.5893 | 3 | * | 0.0278 |
| | 11-deoxycorticosterone | 1.404 | ± | 0.3292 | 3 | 29.43 | ± | 5.611 | 3 | *** | 0.0010 |
| | 17-hydroxyprogesterone | 11.68 | ± | 3.650 | 3 | 3.510 | ± | 0.1374 | 3 | * | 0.0179 |
| | Progesterone | 0.8270 | ± | 0.1931 | 3 | 3.474 | ± | 0.2888 | 3 | *** | 0.0002 |
| **CU-ACC1** | Cortisone | 51.69 | ± | 13.05 | 3 | 26.24 | ± | 1.050 | 3 | * | 0.0281 |
| | Cortisol | 18.83 | ± | 5.901 | 3 | 311.4 | ± | 84.42 | 3 | ** | 0.0039 |
| | Corticosterone | 1.052 | ± | 0.2793 | 3 | 61.10 | ± | 32.75 | 3 | * | 0.0337 |
| | 11-deoxycortisol | 13.67 | ± | 3.38 | 3 | 3.31 | ± | 1.15 | 3 | ** | 0.0074 |
| | Androstenedione | 5.57 | ± | 1.37 | 3 | 1.52 | ± | 0.5 | 3 | ** | 0.0085 |
| | 11-deoxycorticosterone | 0.53 | ± | 0.11 | 3 | 0.36 | ± | 0.18 | 3 | ns | 0.2351 |
| | Testosterone | 1.22 | ± | 0.28 | 3 | 0.18 | ± | 0.06 | 3 | ns | 0.2805 |
| | 17-hydroxyprogesterone | 1.67 | ± | 0.4 | 3 | 1.06 | ± | 0.42 | 3 | ns | 0.1411 |
| | Progesterone | 0.000 | ± | 0.000 | 3 | 0.196 | ± | 0.095 | 3 | * | 0.0235 |
| **MUC1** | 11-deoxycortisol | 0.00 | ± | 0.00 | 3 | 0.81 | ± | 0.69 | 3 | ns | 0.1111 |
| | Androstenedione | 0.20 | ± | 0.03 | 3 | 0.64 | ± | 0.63 | 3 | ns | 0.2953 |
| | 17-hydroxyprogesterone | 0.43 | ± | 0.11 | 3 | 1.30 | ± | 1.39 | 3 | ns | 0.3396 |
| | Cortisone | 0.35 | ± | 0.20 | 3 | 2.96 | ± | 4.09 | 3 | ns | 0.3317 |
| **TVBF7** | 11-deoxycortisol | 0.00 | ± | 0.00 | 3 | 0.29 | ± | 0.51 | 3 | ns | 0.3739 |
| | Androstenedione | 0.04 | ± | 0.02 | 3 | 0.34 | ± | 0.39 | 3 | ns | 0.2578 |
| | 17-hydroxyprogesterone | 0.20 | ± | 0.08 | 3 | 1.36 | ± | 0.77 | 3 | ns | 0.0598 |
| **BCH-ACC3A** | Corticosterone | 111.19 | ± | 35.93 | 4 | 759.57 | ± | 80.84 | 3 | **** | <0.0001 |
| | 11-deoxycortisol | 9.06 | ± | 3.11 | 4 | 57.02 | ± | 5.32 | 3 | **** | <0.0001 |
| | 11-deoxycorticosterone | 7.23 | ± | 2.38 | 4 | 11.39 | ± | 1.87 | 3 | ns | 0.0556 |

**Table S4. LC-MS/MS hormonal output of different ACC cell lines grown in adherent and spheroid culture.**

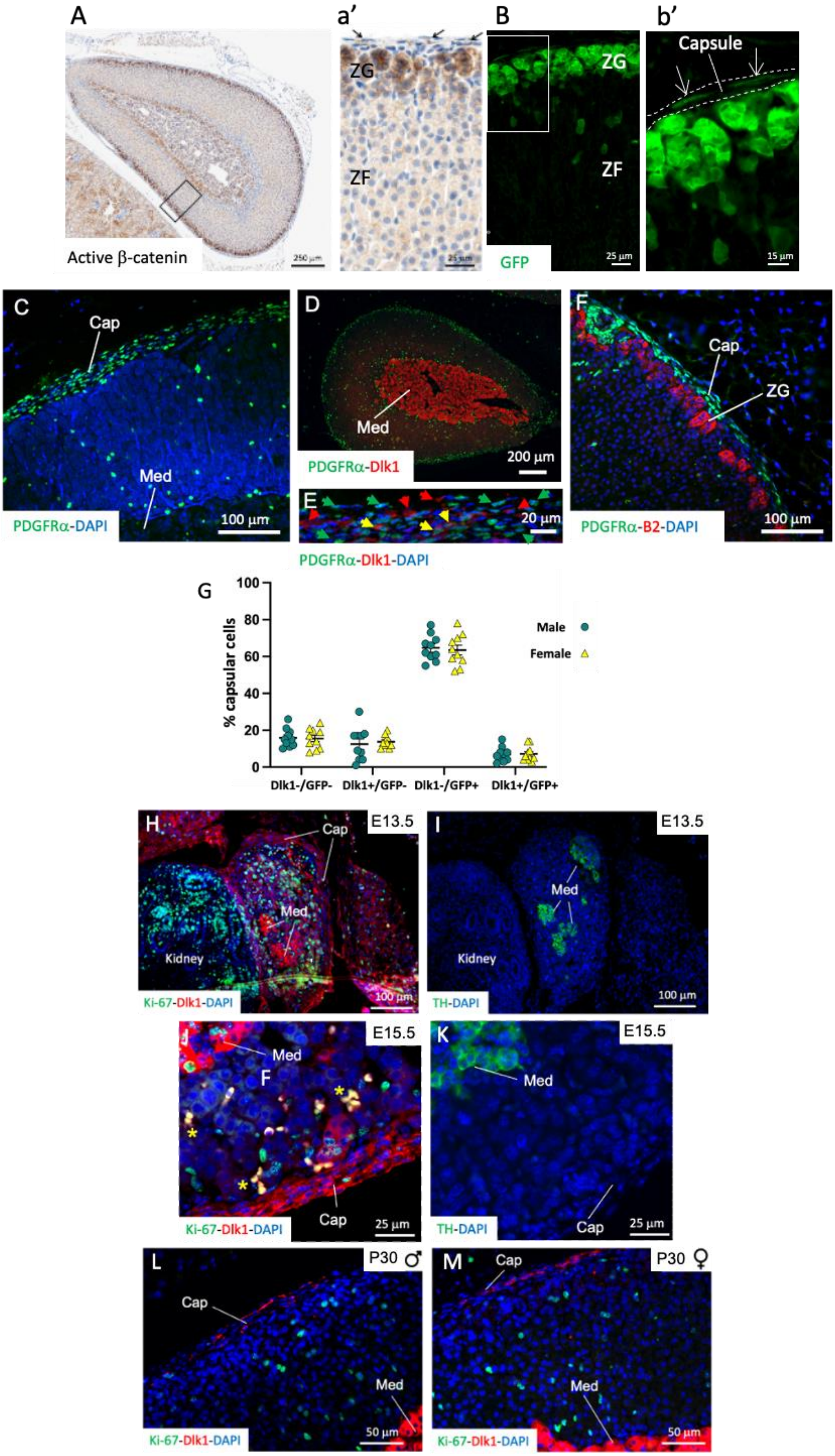
A
Active β-catenin
250 μm
a'
ZG
ZF
25 μm
B
ZG
ZF
GFP
25 μm
b'
Capsule
15 μm
C
Cap
Med
PDGFRα-DAPI
100 μm
D
Med
PDGFRα-Dlk1
200 μm
E
20 μm
PDGFRα-Dlk1-DAPI
F
Cap
ZG
PDGFRα-B2-DAPI
100 μm
G
% capsular cells
Male
Female
Dlk1-/GFP-
Dlk1+/GFP-
Dlk1-/GFP+
Dlk1+/GFP+
H
E13.5
Cap
Med
Kidney
Ki-67-Dlk1-DAPI
100 μm
I
E13.5
Med
Kidney
TH-DAPI
100 μm
J
E15.5
Med
F
Cap
Ki-67-Dlk1-DAPI
25 μm
K
E15.5
Med
Cap
TH-DAPI
25 μm
L
P30 ♂
Cap
Med
Ki-67-Dlk1-DAPI
50 μm
M
P30 ♀
Cap
Med
Ki-67-Dlk1-DAPI
50 μm

**Figure S1.** A-B) Immunohistochemical staining for active β-catenin (A and a') in the mouse adrenal and comparison with GFP signal from *Axin2Cre* mice (B and b', 14 days chase). Arrows in a' and b' indicate positive capsular cells. C-F) Immunofluorescence staining of adrenals obtained from PDGFR$\alpha^{EGFP}$ mice and stained for GFP (indicating PDGFR$\alpha^{EGFP}$ cells, C), GFP and Dlk1 (D and E), GFP and Cyp11b2 (B2, F). G) Percentage of cells expressing or not expressing Dlk1 and PDGFR$\alpha^{EGFP}$ in male and female adrenals. H-M) Immunofluorescence staining for Ki-67, Dlk1 and DAPI at E13.5 (H), E15.5 (J) and at P30 (L, M) showing occasional colocalization of Ki-67 with Dlk1 in the capsule and subcapsular clusters during development and never postnatally. Developing medulla was localized with Tyrosine Hydroxylase (TH) antibodies (I and K) in adjacent sections. Yellow asterisks in panel F indicate auto fluorescent erythrocytes. Cap, capsule; Med, medulla; ZG, Zona Glomerulosa; ZF, Zona Fasciculata. Nuclei in all panels except B are stained with DAPI, or with hematoxylin (A).

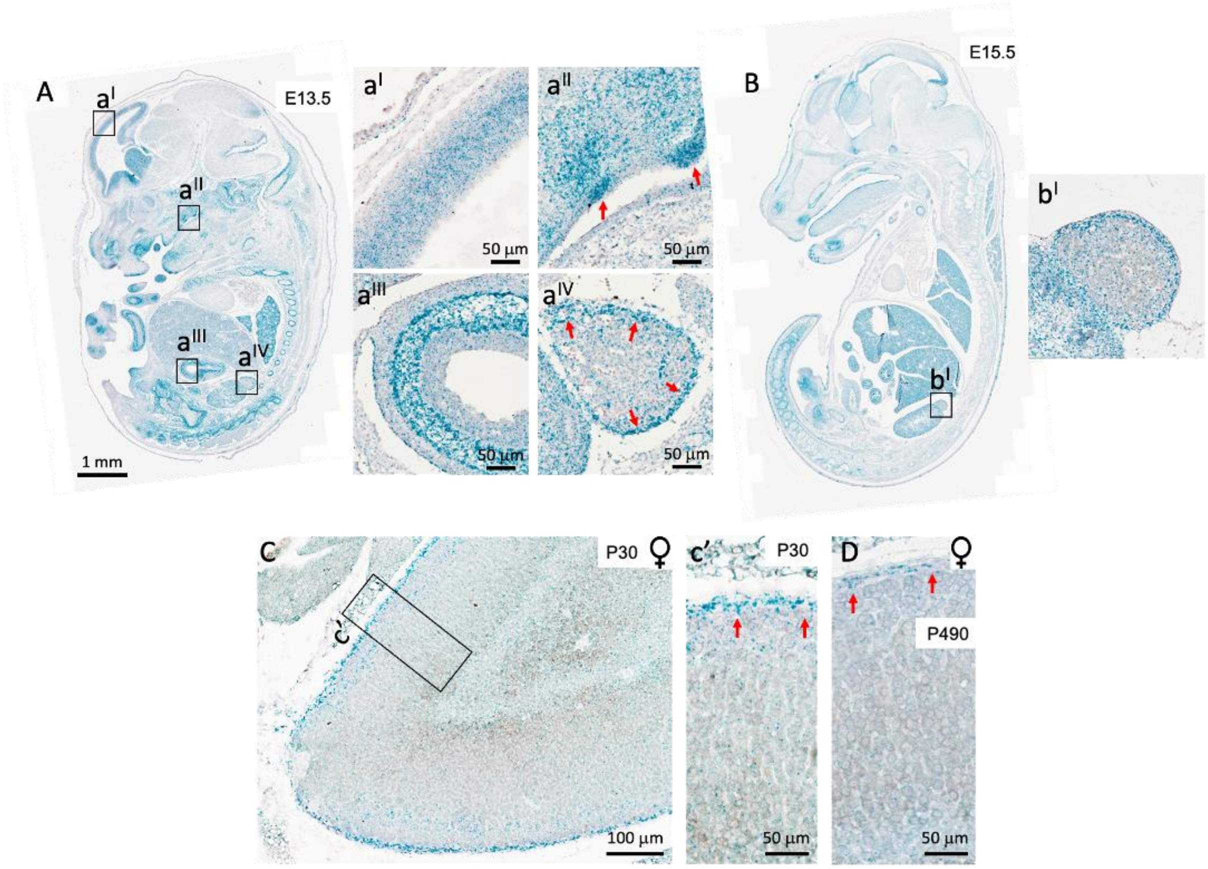


**Figure S2. *Gli1* maintains a strong capsular expression throughout the lifetime in mice.** RNAScope detection of *Gli1* mRNA at E13.5 (A), E15.5 (B), P30 (C) and P490 (D) showing strong staining in the capsule during development throughout old age. Expression of *Gli1* in male adrenals is similar to females (not shown). Known areas of *Gli1* expression are shown in panels a' (neocortex), a'' (mesenchyme by the papillae), and a''' (mesenchyme by the stomach).

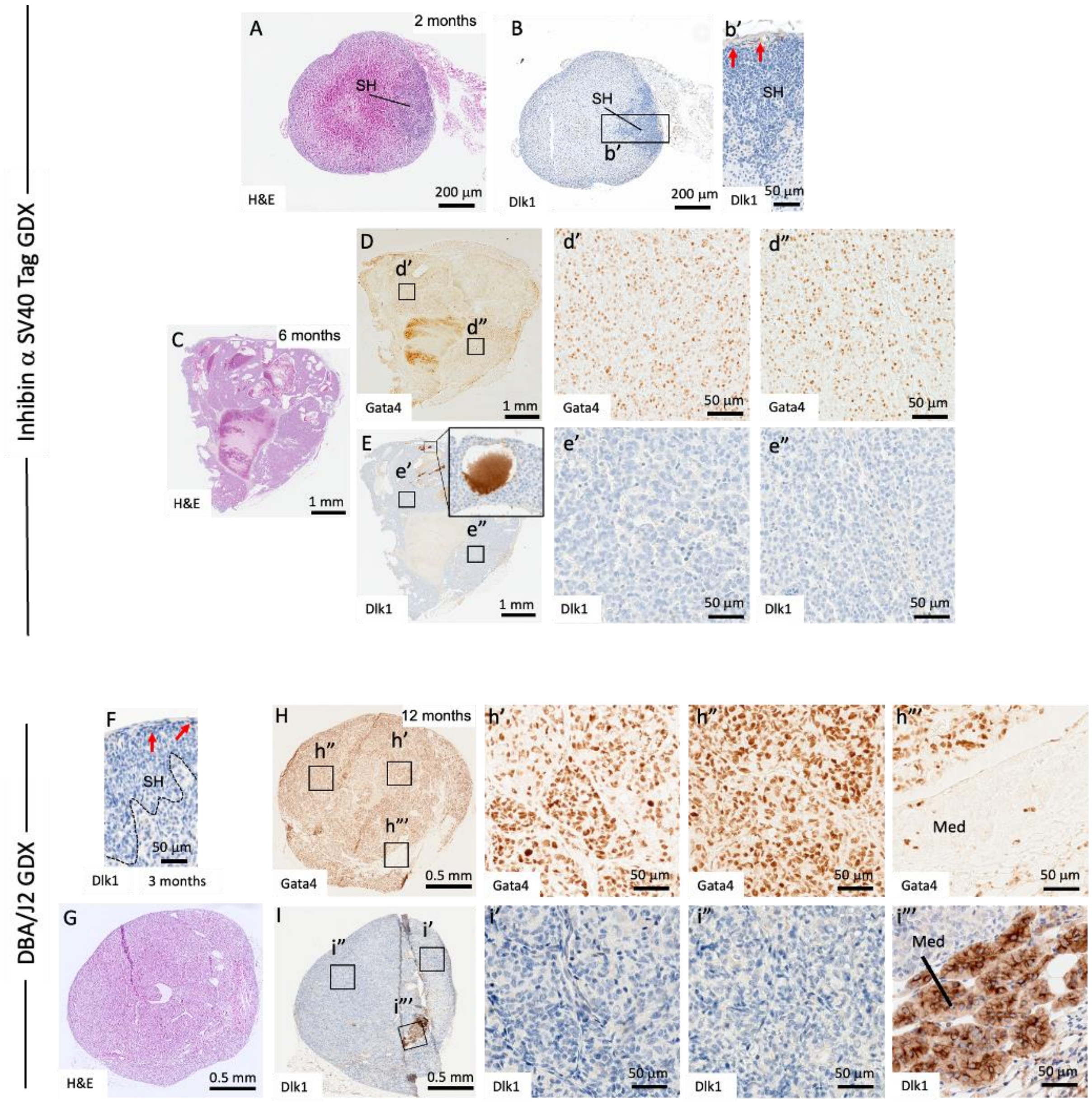


**Figure S3. Dlk1 is not expressed in adrenal subcapsular hyperplasia (SH) and subsequent adrenocortical tumors** A-E) Inhibin α SV40 Tag mice underwent prepubertal gonadectomy (GDX), resulting in the development of SH at 2 months (A and B), followed by adrenocortical tumors at 6 months (C-E). Expression of Dlk1 was restricted to the external capsule in SH (red arrows in b') and was not expressed in tumors (E). Tumors expressed Gata4 transcription factor (D). Magnified panel in E shows artefactual staining in luminal secretions. F-I) DBA/2Jmice underwent prepubertal gonadectomy (GDX), resulting in the development of SH at 3 months (F), followed by adrenocortical tumors at 12 months (G-I). Expression of Dlk1 was restricted to the external capsule in SH (red arrows in F) and was not expressed in tumors (I), except in the remnant of the medulla (Med, panel i'''). Tumors expressed Gata4 transcription factor (H).

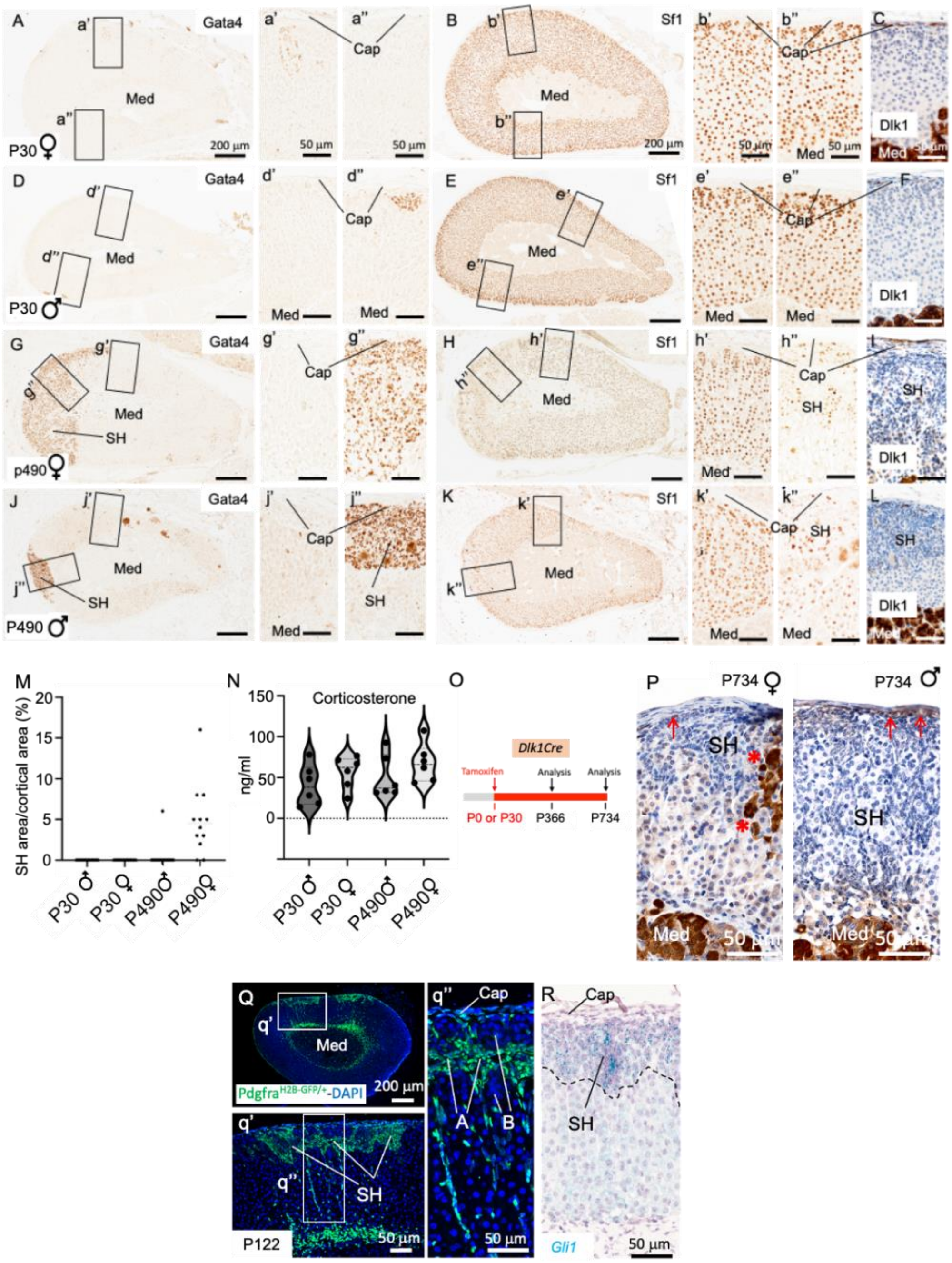

**Figure S4. SH foci in aged mice are not enriched in nor derived from Dlk1 cells.** A-L) Expression of Gata4, Sf1, and Dlk1 in young (P30) and aged (P490) adrenals. Large SH foci were observed in 11/12 females and in 1/12 males. Panels J and K are not representative of male adrenals but show the only male which developed SH. Dlk1 expression (panels C, F, I and L) was restricted to the outermost part of each SH, while both A and B cells were negative for Dlk1 expression. M) % area of SH, expressed as the ratio of Gata4$^{+}$ cells vs total cortical area. N: LC-MS/MS levels of corticosterone in sera of P30 and P490 mice. O) Schematic of tamoxifen induction in *Dlk1Cre* mice. P) A total of 57 SH foci from 24 mice (3 of each males, females, injection at P0, P30, chase up to P366 and P734) were analyzed for RFP expression; none of the SH were RFP positive, with RFP cells confined to the outermost part of the SH (red arrows), in the medulla, and in the occasional non-SH cluster/columns representing Dlk1 cortical progeny (red asterisks in P). Q-R) SH A cells are positive for PDGFRα and *Gli1* expression. Cap, capsule; Med, medulla.

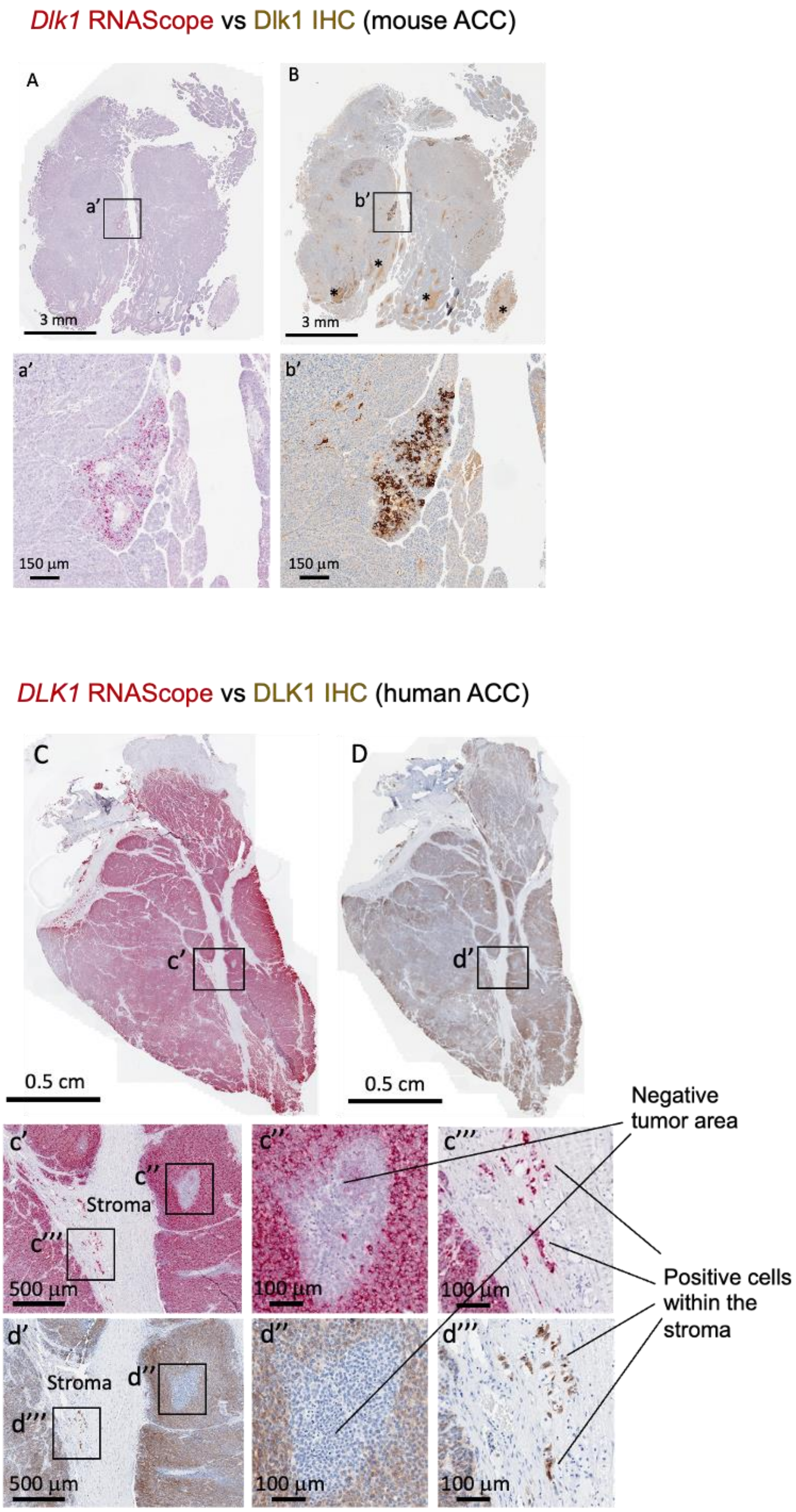


**Figure S5. The expression profiles of *Dlk1* mRNA and Dlk1 protein in mouse and human ACCs are specular**. Examples of Dlk1 expression in a low (mouse) and high (human) H-score ACC. Consecutive sections of FFPE mouse (*BPCre*) and human ACC were processed for Dlk1 RNAScope (A and C) or IHC (B and D). Asterisks in B indicate IHC artefactual staining in luminal secretions.

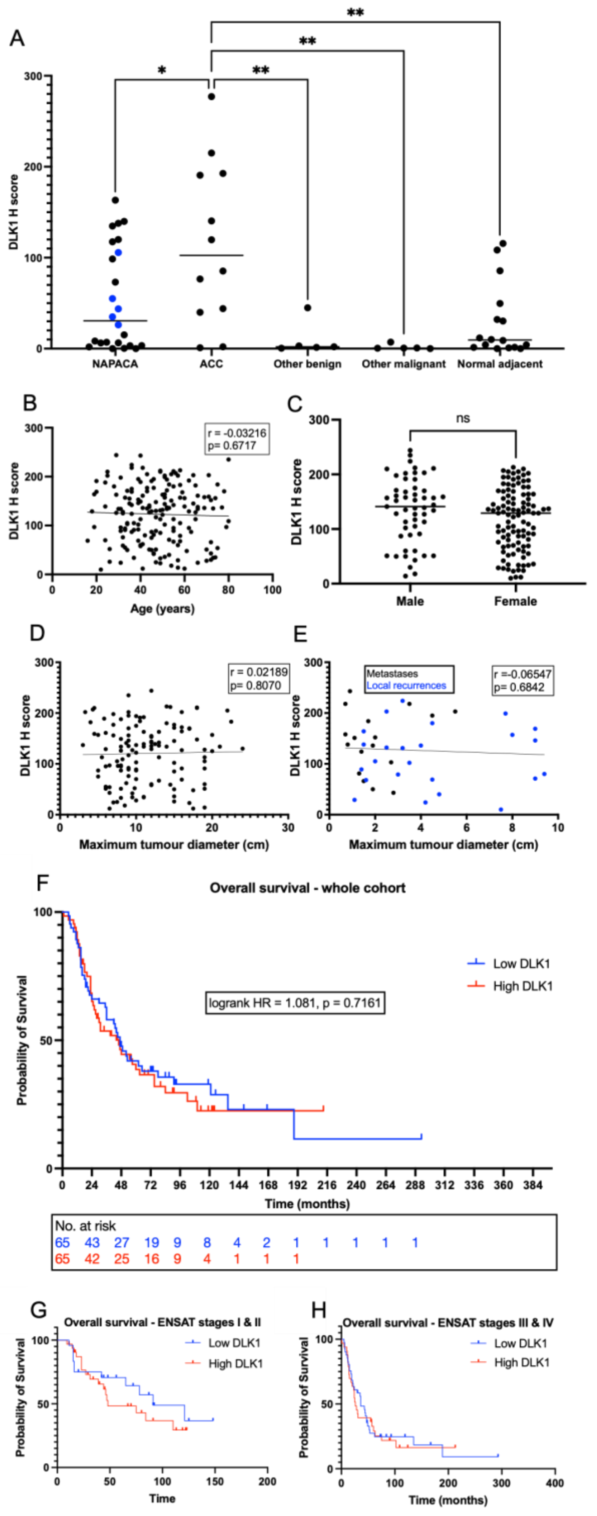


**Figure S6. DLK1 expression is higher in ACC, is not affected by age, sex or tumor size and is not associated with a change in overall survival.**

A) DLK1 expression is higher in ACC than other adrenal pathologies (*London cohort*). B-H) *Würzburg cohort*. DLK1 expression in ACC does not correlate with age (B), sex (C), tumor size in either primary (D) or secondary (E) tumor samples. F) Kaplan Meier curve showing no difference in overall survival with DLK1 expression in entire cohort. G-H) A trend to a survival benefit is seen in ENSAT stage I & II disease (G) but not in stage III & IV disease (H).

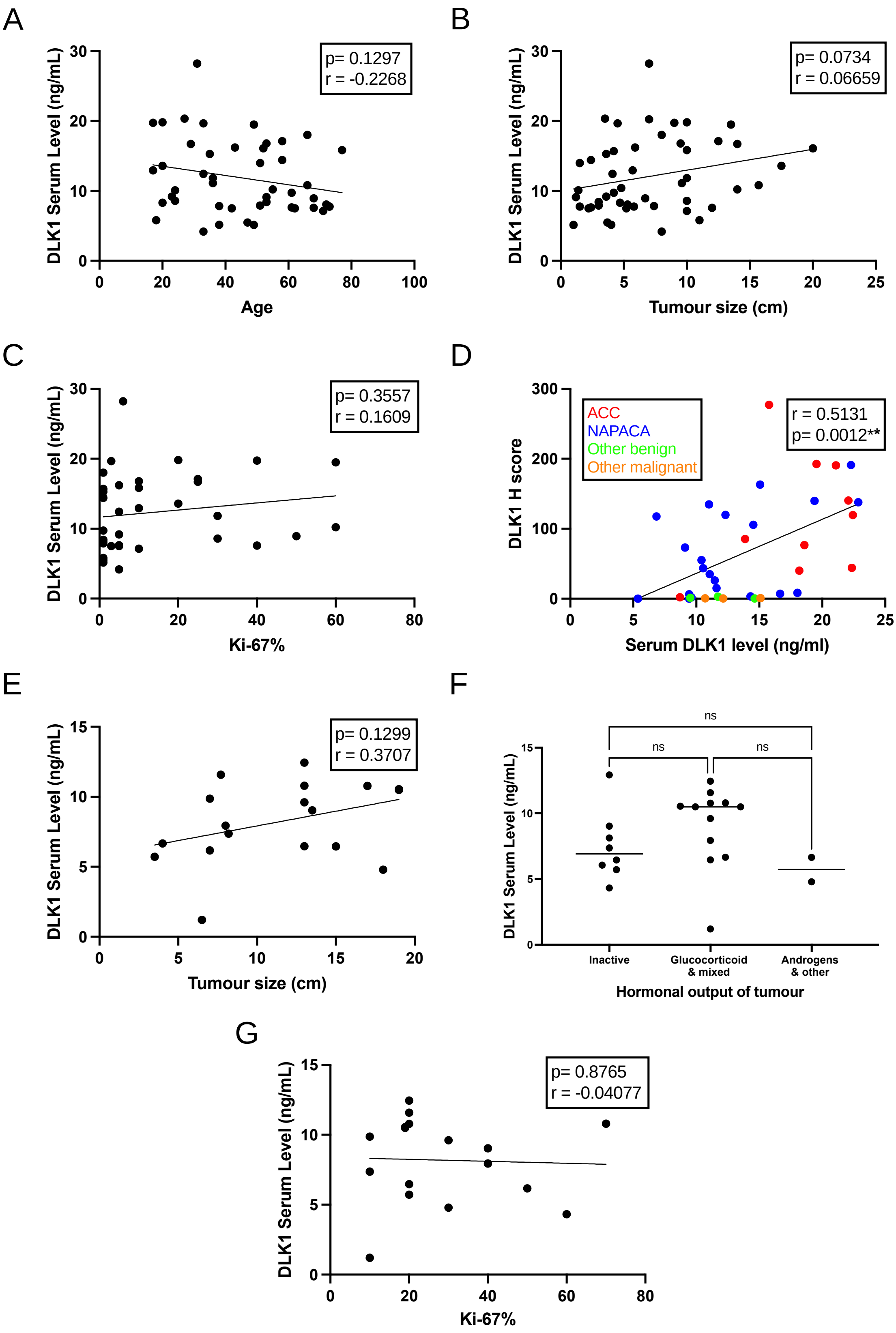


**Figure S7. DLK1 serum levels are not affected by presentation of disease**. A-D) *London cohort.* DLK1 serum levels are not correlated with patient age (A), tumor size (B) or Ki-67% (C). D) DLK1 serum levels correlate with tissue expression in same patients across the entire

London cohort. E-G) *Würzburg cohort*. In ACC alone, DLK1 does not correlate with tumor size (E), tumor hormonal output (F) or Ki-67% (G).

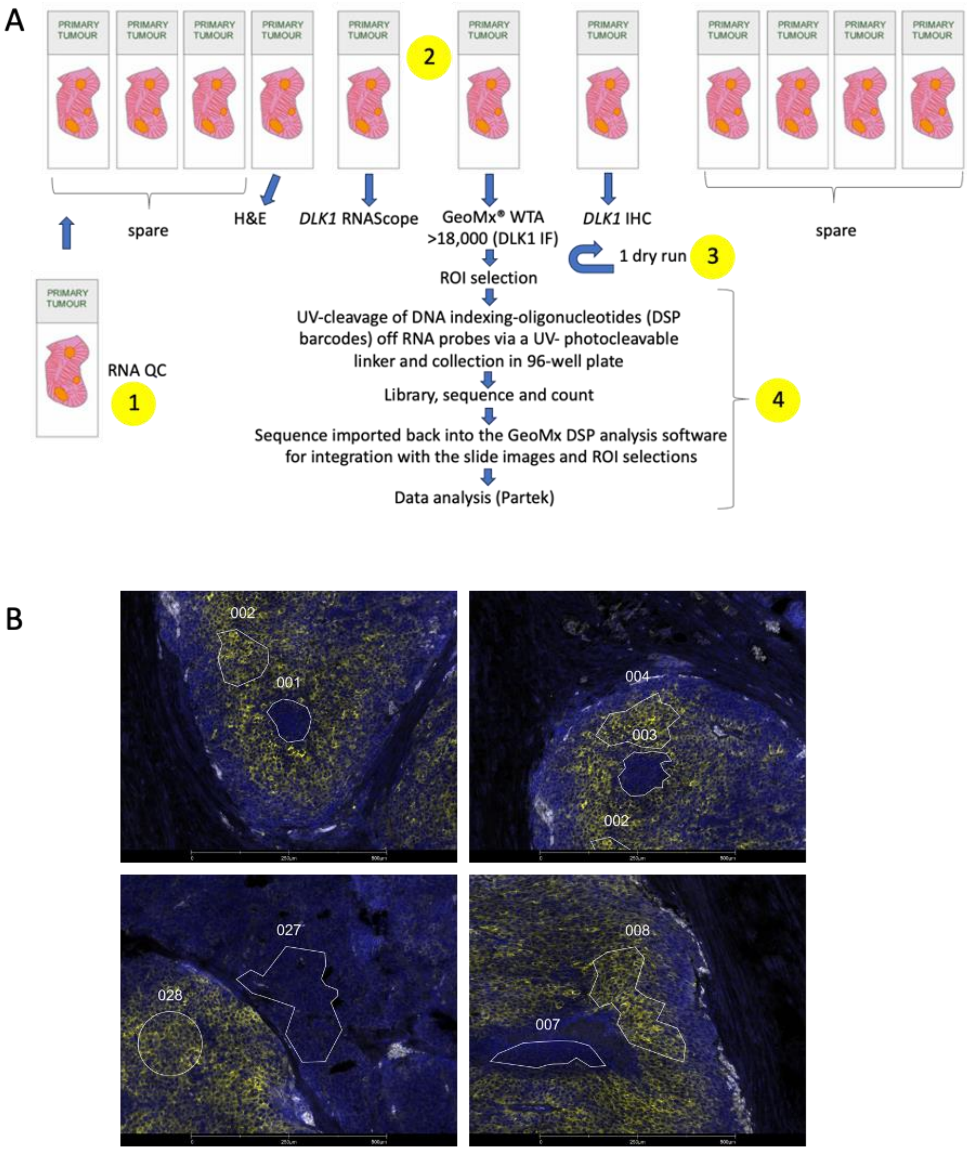


**Figure S8. Workflow of GeoMx Spatial transcriptomics**. A) Consecutive FFPE sections were mounted on to Superfrost Plus slides and two random sections/ACC were processed for mRNA Quality Control (QC) (step 1). After QC, consecutive sections were processed for Hematoxylin and Eosin, DLK*1* RNAScope, DLK1 immunohistochemistry (IHC), DLK1 immunofluorescence (IF), and DLK1 IF according to the GeoMx protocol (step 2). A dry run was done for each sample before the experiment to assess that DLK1 IHC/IF/RNAScope signals were identical to that obtained with the IF from the GeoMx protocol (step 3). After hybridization with Whole Transcriptome Atlas (WTA), Regions of Interest (ROI) were selected based on DLK1 signal, and Digital Spatial Profile (DSP) barcodes collected into 96-well plates (1 ROI/well) via UV-cleavage and microcapillary aspiration, before sequencing the library (Illumina) (step 4). B) Examples of ROI from ACCs based on DLK1 expression (ROI 1, 3, 7, 27 DLK1$^{-}$, ROI 2, 4 ,8 28 DLK1$^{+}$). Stroma and vasculature were excluded.

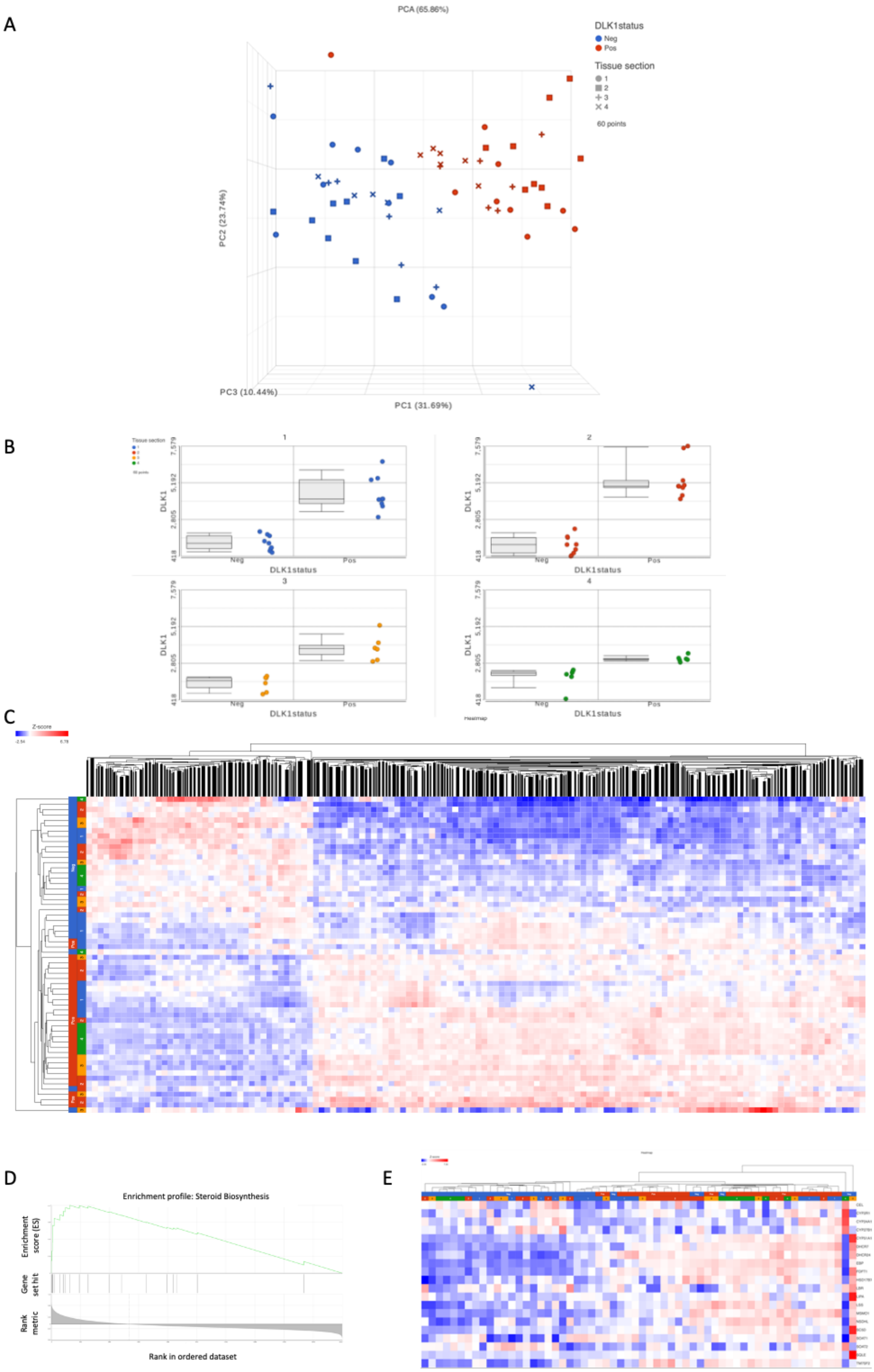
A
PCA (65.86%)
DLK1status
Neg
Pos
Tissue section
1
2
3
4
60 points
PC2 (23.74%)
PC3 (10.44%)
PC1 (31.69%)
B
1
2
3
4
DLK1
7.579
5.192
2.805
418
Neg
Pos
DLK1status
C
Z-score
D
Enrichment profile: Steroid Biosynthesis
Enrichment score (ES)
Gene set hit
Rank metric
Rank in ordered dataset
E

**Figure S9**. **DLK1$^{+}$ cells are endowed with both enhanced steroidogenic potential and clonogenicity.** A) Principal component analysis (PCA) plot of the 4 ACC samples analyzed by spatial transcriptomics, showing the clustering of DLK1$^{+}$ and DLK1$^{-}$ tissue areas. B) *DLK1* mRNA count in the ROI from four ACC samples which were expressing low/nil levels (Neg) or were positive (Pos) for DLK1 protein expression. C) Heatmap showing unsupervised clustering of differentially expressed genes between DLK1$^{+}$ and DLK1$^{-}$ tumor areas. The most upregulated pathway was of steroid biosynthesis (enrichment analysis (D)). E) Heatmap showing unsupervised clustering of differentially expressed genes in steroid pathway dataset.

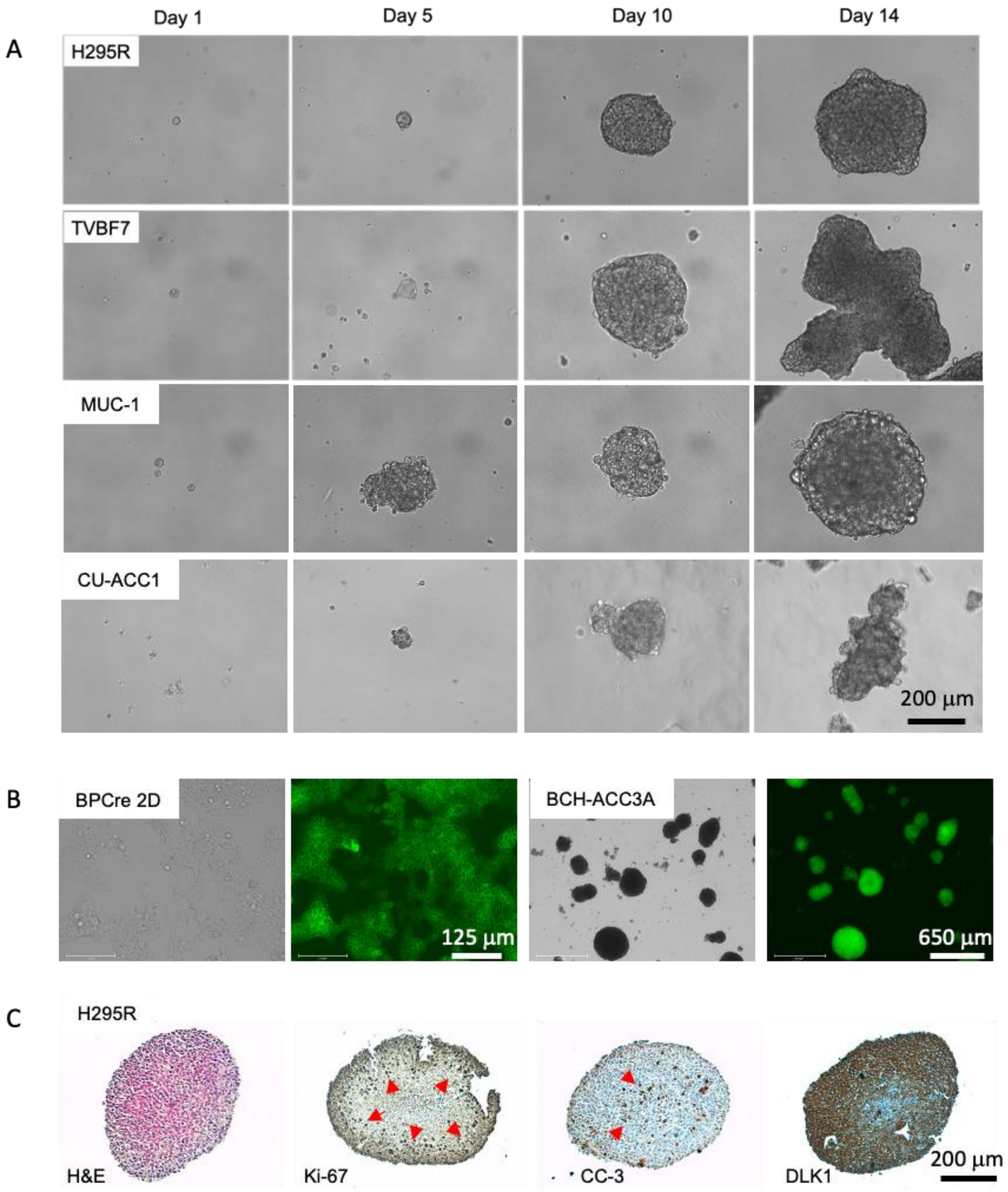


**Figure S10**. **Derivation of 3D spheroid cultures from human and mouse ACC lines.** A) Time course of spheroids generation in H295R, TVBF7, MUC-1 and CU-ACC1 human ACC cells. B) Bright field and fluorescent images of BCH-ACC3A ACC mouse cells grown in adherent (2D) and as spheroids (3D). C) Paraffin sections from PFA-fixed H295R spheroids were stained with H&E, Ki-67, Cleaved caspase 3 (CC-3) and Dlk1. The majority of Ki-67$^+$ cells are located at the periphery, while signal for Cleaved caspase 3 is mainly in the center of the spheroid, indicating normoxic and hypoxic areas, respectively. Note the strong expression of DLK1.
